# Oscillations without cortex: Working memory modulates brainwaves in the endbrain of crows

**DOI:** 10.1101/2022.02.01.478165

**Authors:** Lukas Alexander Hahn, Dmitry Balakhonov, Mikael Lundqvist, Andreas Nieder, Jonas Rose

## Abstract

Complex cognition requires coordinated neuronal activity at the network level. In mammals, this coordination results in distinct dynamics of local field potentials (LFP) that have been central in many models of higher cognition. Because these models are based on mammalian data, they often implicitly assume a cortical organization. Higher associative regions of the brains of birds do not have cortical layering, yet these regions have neuronal single-cell correlates of higher cognition that are very similar to those found in mammals. Here we recorded LFP in the avian equivalent of prefrontal cortex while crows performed a highly controlled and cognitively demanding working memory task, adapted from monkeys. To further ensure that recordings reflected only cognitive processes detached from motor-related activities we trained and monitored the animals to keep their head still. We found signatures in local field potentials, modulated by working memory. Frequencies of a narrow gamma (30-59 Hz) and the beta band (13-19 Hz) contained information about the location of the target items on the screen and were modulated by working memory load. This indicates a critical involvement of these bands in ongoing cognitive processing. We also observed bursts in the beta and gamma frequencies, similar to those observed in monkeys. Such bursts are a vital part of ‘activity silent’ models of working memory. Thus, despite the lack of a cortical organization the avian associative pallium can create LFP signatures reminiscent of those observed in primates. This points towards a critical cognitive function of oscillatory dynamics evolved through convergence in species capable of complex cognition.

**Relevance statement:** Contemporary models of higher cognition, like those of working memory, often include temporal dynamics of neural activity such as gamma oscillations. Birds and mammals convergently evolved these cognitive functions and here we show that, despite the large evolutionary distance and largely different brain organization, crows share many of the oscillatory fingerprints reported in primates. This indicates that neural networks required for such LFP phenomena have evolved in parallel and may be critical to higher cognition.

## Introduction

To perform the computations underlying complex cognition, the neuronal ensembles of our brains must be coordinated, otherwise, the chatter of a billion neurons may produce only noise (Lisman, 1997; Miller et al., 2018; Naud & Sprekeler, 2018). Notably, the spiking of individual neurons follows a tight temporal organization that results in regular patterns of excitation and inhibition. At the network level, these patterns of activity can be observed in fluctuations of electrical local field potentials (LFP) that oscillate at different frequencies (Buzsáki et al., 2012, 2013; Buzsáki & Wang, 2012). These frequencies are commonly clustered into bands, for example, the gamma band of frequencies above 30 Hz. Gamma oscillations are likely generated in the superficial layers of cortex (Bastos et al., 2018; Buffalo et al., 2011; Maier et al., 2010), from perisomatic currents around the similarly oriented pyramidal cell layer and they arise from feedback inhibition between pyramidal cells and somatic targeting parvalbumin-positive inhibitory neurons (Buzsáki et al., 2012; Buzsáki & Wang, 2012; Cardin et al., 2009; Carlén et al., 2012; Traub et al., 1996). Functionally, the gamma band has been suggested to be relevant for inter-regional communication of neuronal populations (Fries, 2015), and to play a key role in executive control (Miller et al., 2018). Thus, understanding these coordinated computations is the key to unlocking a functional model of higher cognition.

A cornerstone of complex cognition is working memory (WM), which enables an animal to actively retain and manipulate a limited amount of information to guide behavior (Baddeley et al., 2021). WM is also particularly well suited to investigate higher cognition from a comparative perspective. It was described almost simultaneously in humans and pigeons (Baddeley & Hitch, 1974; Honig, 1978). Furthermore, birds and mammals show similar WM performance (Balakhonov & Rose, 2017; Gibson et al., 2011). For example, the capacity of WM, the number of individual items that can be maintained simultaneously, is comparable between crows and macaque monkeys (Balakhonov & Rose, 2017). Even single neuron correlates of WM in birds are virtually identical to those in mammals (Ditz & Nieder, 2016, 2020; Moll & Nieder, 2015; Rinnert et al., 2019; Rose & Colombo, 2005) and we recently found that this also extends to the neurophysiological limits of WM capacity (Buschman et al., 2011; Hahn et al., 2021).

Given the large evolutionary distance between the species, these similarities are likely the result of convergent evolution (Emery & Clayton, 2004; Güntürkün & Bugnyar, 2016) and they are sharply contrasted by prominent anatomical differences. Most notably, birds lack the mammalian separation between grey and white matter along with the highly structured organization of the neocortex (Güntürkün & Bugnyar, 2016; Harris & Shepherd, 2015). While recent data suggest a cortex-like circuitry in sensory regions of the avian pallium, a layered neocortex-like structure is absent in associative avian brain regions that are crucial to WM function (Stacho et al., 2020). This includes the avian equivalent of PFC, the nidopallium caudolaterale (NCL), which shares many defining properties of the PFC, including the dense dopaminergic innervation, multimodal sensory afferents, premotor projections, and neuronal correlates for WM (Güntürkün & Bugnyar, 2016; Herold et al., 2011; Kröner & Güntürkün, 1999; Nieder, 2017; Waldmann & Güntürkün, 1993).

Modern models of WM are heavily influenced by the observation of temporal dynamics in the mammalian PFC. In particular, gamma oscillations are closely associated with WM-related processes (Howard et al., 2003; Kornblith et al., 2016; Lundqvist et al., 2016; Roux et al., 2012; Tallon-Baudry et al., 1998). The highly structured organization of the layered mammalian neocortex is an ideal substrate to generate and investigate such oscillations (Einevoll et al., 2013). Consequently, models of temporal dynamics are almost exclusively built on mammalian data. However, whether these cognitive oscillations *require* the specific layered organization of the cortex is unclear. It has even been argued that oscillations could be an epiphenomenon of the underlying network architecture rather than a functional process in itself (Merker, 2013; Ray & Maunsell, 2015). Therefore, the investigation of LFP in avian associative brain regions, lacking the layered organization of the cortex, offers a unique comparative perspective.

To date, only relatively few studies have investigated modulations of LFP in birds. Most prominently the optic tectum and neighboring tegmental nuclei show modulation in the gamma range during attention (Goddard et al., 2012; Neuenschwander & Varela, 1993; Sridharan et al., 2011; Sridharan & Knudsen, 2015). Gamma band modulations were further reported in the avian forebrain during birdsong (Brown et al., 2021; Lewandowski & Schmidt, 2011; Spool et al., 2021), and in the avian hippocampal formation *in vitro* (Dheerendra et al., 2018) and during sleep (van der Meij et al., 2020). However, these observations cannot answer the question of whether oscillations underlie higher cognition since they were either made in the neatly layered optic tectum, were tightly linked to motor behavior, or occurred in sleeping birds.

Thus, descriptions of oscillatory dynamics in the non-layered endbrain of birds that are tied to abstract cognition such as WM are still lacking. Hence, it remains unknown if the single-cell similarities extend to oscillatory population dynamics that underlie higher cognition in mammals, or if birds have such cognition without oscillations. If they existed and played comparable roles in avian and mammalian WM, it would be valuable evidence towards general, cross-species mechanisms supporting higher-order cognition.

## Results

To investigate LFP dynamics in the avian brain during a complex form of cognition, we trained crows on a multi-item working memory task, previously used for probing WM capacity in crowns and primates (Balakhonov & Rose, 2017; Buschman et al., 2011). On each trial, the crows were presented with a variable number of colored squares that they had to retain over a memory delay. Subsequently, the colors reappeared and the birds indicated with a single peck which of the squares now had a different color (Fig. 1A). The performance of the crows was load-dependent, gradually declining with higher loads. Median performances for item loads (ipsilateral to change) of one, two, and three were 95.88 %, 78.31 %, and 58.21 %, respectively. This result is very similar to the performance reported in monkeys in the same task (Buschman et al., 2011), and has been discussed in detail in a previous study (Balakhonov & Rose, 2017).

**Figure 1:**
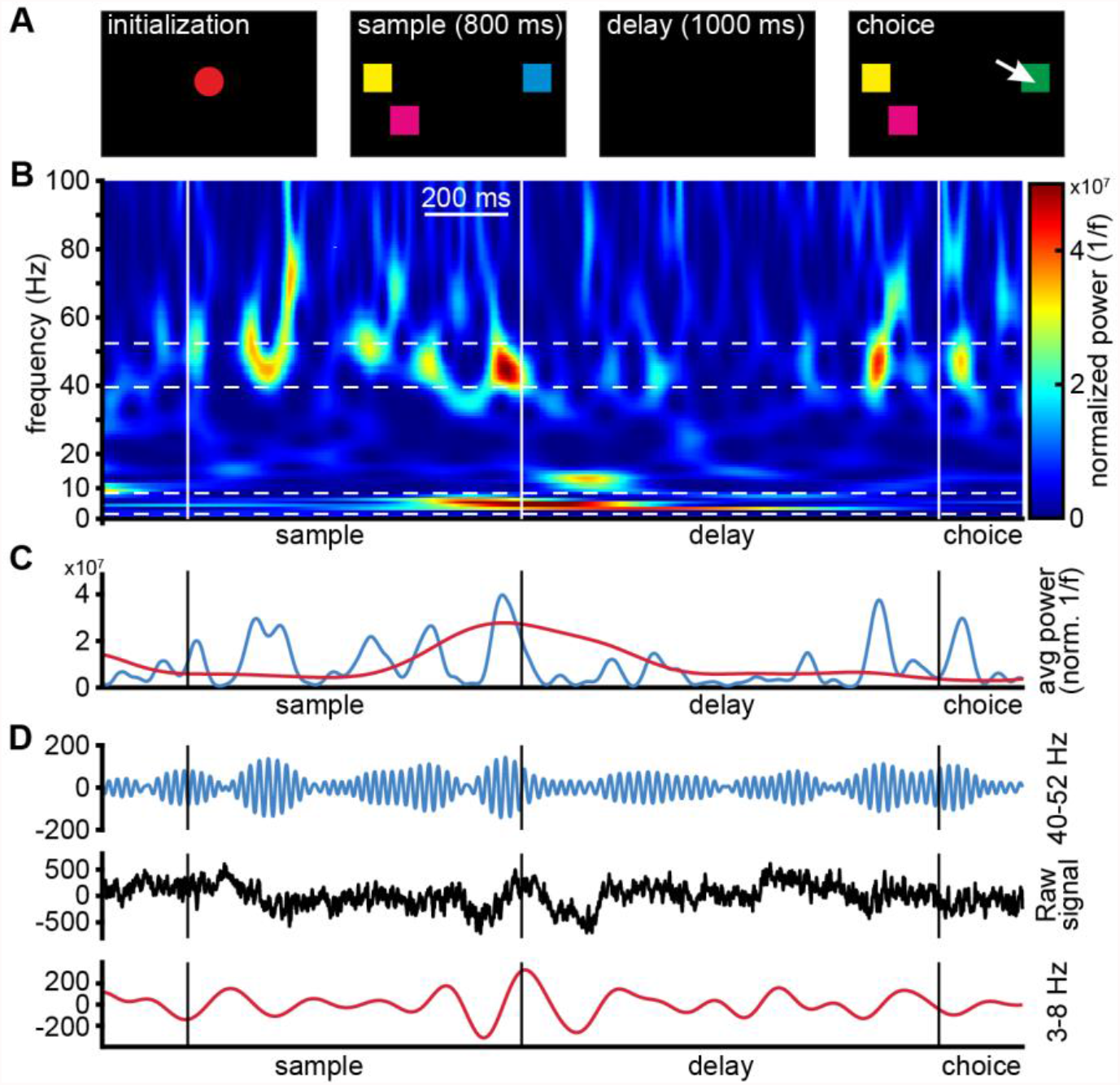
(**A**) Behavioral protocol. After the bird initiated a trial by acquiring and holding head fixation, the sample stimuli (2-5 colored squares distributed so that 0-3 colored squares appeared on each half of the screen) were presented. Birds retained head fixation and maintained color information over a memory delay, until the choice stimuli were presented (identical in color and location to those of the sample phase, except for one square that had changed color). Birds then indicated the square that changed color between sample and choice by pecking on it. (**B**) Single-trial example of time-frequency power of LFP. Power was elevated during the transition from sample to delay phase in a band between 3 and 8 Hz. Higher frequencies between 40 and 52 Hz showed recurring increases of power in short bursts during the sample and the delay period, notably also towards the end of the delay. (**C**) Mean power of the selected bands across time (3-8 Hz, red, and 40–52 Hz, blue). The visible peaks correspond to the warmer colors in panel B. (**D**) Raw unfiltered LFP signal (black), and the same signal, band-pass filtered in the range of higher frequencies (blue), and of the lower range frequencies (red). The respective frequency components of the raw signal become visible as their amplitude increased and decreased over time.

### LFP in the endbrain of crows is task modulated

To investigate if WM modulates oscillations in a comparable way in crows as in primates, we analyzed LFP recorded throughout NCL from a total of 336 electrodes. We performed spectral decomposition of the recorded signal using Morlet-wavelet convolution, after removing neuronal spiking artifacts and 50 Hz line noise (see Methods for details). LFP power was affected throughout the time course of a trial in a frequency-dependent manner. To facilitate the comparison to results obtained in primates we subdivided frequencies into the commonly recognized LFP bands, i.e., ‘theta’, ‘alpha’, ‘beta’, and ‘gamma’ (Miller et al., 2018).

We observed modulation of LFP power in a narrow gamma frequency band (40-52 Hz) during the sample phase and delay phase, as well as high levels of power in a 3-8 Hz frequency band toward the end of the sample phase (Fig. 1B and C). This was also observed in the raw signal trace, most prominently in the sample and towards the end of the delay, when the individual frequency components contributed most to the composite signal (indicated by higher frequency amplitudes in Fig. 1D).

Were specific frequency bands consistently affected by the ongoing cognitive task? We tested trial averaged LFP power during the trial against stable baseline power (see Methods for details). The observations, made at the single-trial level, were consistent across trials (Fig. 2, example electrode). Power in the low band was significantly suppressed during the early sample, and at the end of the delay (Fig. 2, bottom). The high-frequency band (gamma) was significantly elevated relative to baseline during the late sample and towards the end of the delay phase (Fig. 2, top; see SFig. 1 for statistical results and further details).

**Figure 2:**
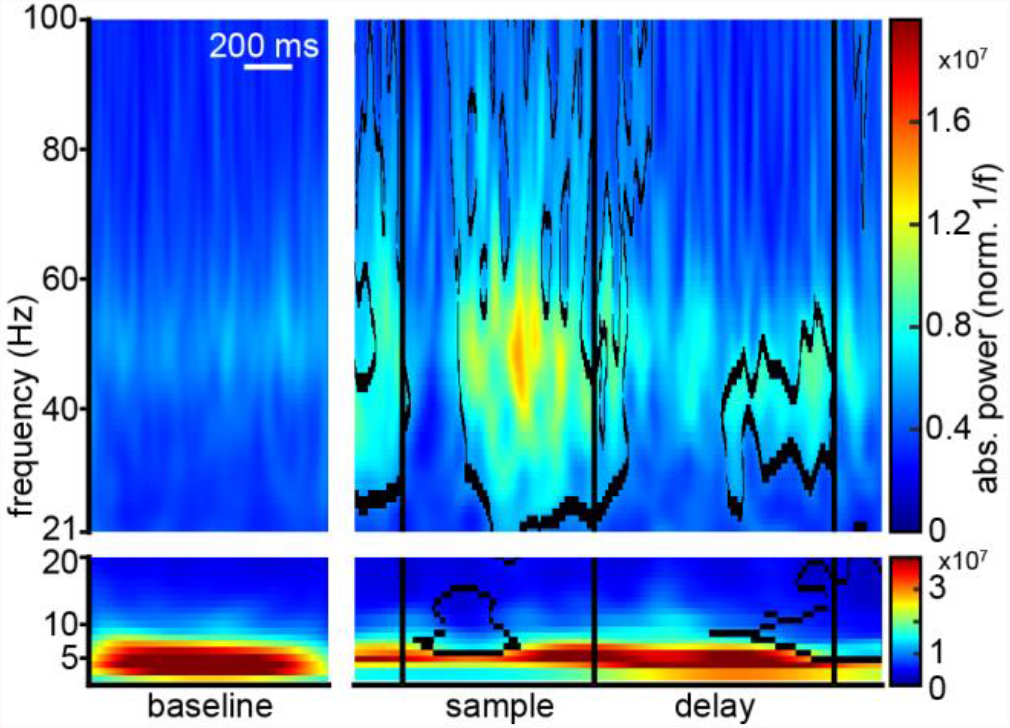
Average time-frequency power of LFP of a single electrode of a single session during the baseline period (1 second during the middle of the inter-trial-interval) and during the trial period. The duration of the pre-sample period was variable dependent on behavior, it could therefore not be used as baseline, and it contains motion and stimulus-viewing. In the sample phase an increase in gamma power, and a decrease in alpha/beta power is detectable. Outlined areas indicate power values significantly different from baseline. Higher and lower frequencies were split to highlight their respective power range that scales with 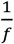.

This shows that modulations of LFP were generated in NCL in narrow and well-defined frequency bands. These modulations reflected processing in the different task phases, and were not motor-related, as the birds had to retain a stable head up until the choice. Because the gamma frequency range was most affected by our task, we focused on electrodes that showed modulations in that range. We examined the overall modulations of recorded LFP power from all electrodes with significant gamma modulation (see Supplementary section 2 for more details).

The sampled average signal showed that the task phases strongly affected the LFP. Both low gamma frequencies (33-48 Hz ‘low gamma’) and beta band frequencies (13-19 Hz ‘beta’) showed a distinct modulation by the task. The low gamma band was shortly suppressed after the sample onset, followed by an increase in power towards the end of the sample phase (Fig. 3A top). In the memory delay phase power of these frequencies remained at an elevated level (relative to baseline), and ramped up towards the end of the delay leading up to the choice. Beta frequencies initially showed strong suppression of power during the early sample phase (Fig. 3A bottom) and returned to baseline levels toward the late sample and early delay. Power was again suppressed towards the end of the delay phase, leading up to the choice.

**Figure 3:**
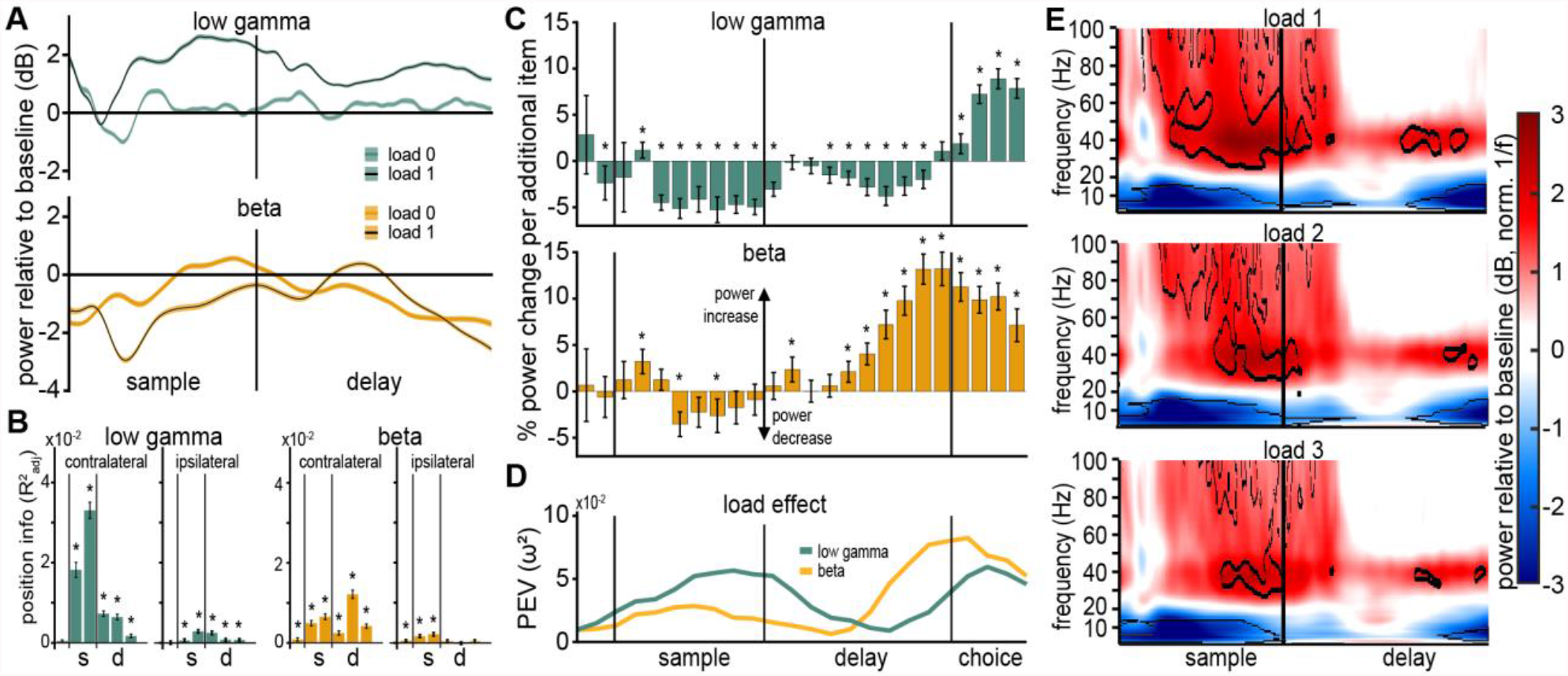
(**A**) LFP in the gamma (top) and beta (bottom) are modulated by working memory. At load 0 no stimuli were presented contralateral to the electrode, at load 1 a single contralateral stimulus was presented during the sample period. (**B**) Position information 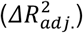 contained in average power of the low gamma and beta band (400 ms bins). Power of the low gamma and of the beta frequency band contained information about the contralateral positions of stimuli, in contrast information about the positions of the ipsilateral stimuli was much smaller. Position information for low gamma frequencies was more pronounced during the sample phase than during the delay phase. Stars indicate significance at the Bonferoni corrected alpha level (α = 0.0083; refer to SFig. 3 for other frequency bands). (**C**) Average change in power per added item (100 ms bins). The low gamma frequency band (32-47 Hz) shows a reduction of power with every added item throughout the sample delay phase, but gains power with every added item in the choice phase. The beta frequency band (12-29 Hz) shows a consistent increase in power with every added item throughout sample and delay phase, notably peaking towards the end of the delay. (**D**) Quantification of the load effect depicted in (A), as percent explained variance by factor power (ω^2^). (**E**) WM load affected the time-frequency power of LFP. Average power of all electrodes with significant gamma band modulation, relative to baseline (in decibel), for load 1-3. Lower frequencies show a general suppression of power, relative to baseline, while higher frequencies show a general increase in power. The tree panels depict different WM-load (number of items contralateral to the recording electrode). Outlined areas indicate significant differences from baseline.

### Gamma modulation reflected working memory processing

The described modulations in power have so far been linked to the processing of the WM task, divided into the processing of presented memory items (during sample), their maintenance (during the delay), and in anticipation of the upcoming change detection (towards the end of the delay). We investigated if our WM task caused further modulation of LFP that reflected cognitive processing of relevant stimulus dimensions, by analyzing if power of these bands contained information about the location of the presented items and if the number of items affected power.

To estimate information about ipsilateral and contralateral positions, we applied the method of (Kornblith et al., 2016), performing model comparisons of generalized linear models (see Methods for details). For all electrodes that had significant gamma modulation (see Supplementary section 2), we derived position information, needed to solve the task, for the ipsilateral and contralateral locations by quantifying the difference of model fits 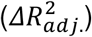, in six 400 ms intervals (pre-sample; early/late sample; early/mid/ late delay).

In general, power contained information about the (task relevant) locations of presented squares (Fig. 3B). This information was most prominently present during the sample phase. We found that low gamma power had significant position information during the sample for the contralateral side of the screen (early and late sample, mean (± SEM): 0.0182 (± 0.0019), F(1,1247) = 379.83, p < 0.0001, ω^2^ = 0.2327 and 0.0330 (± 0.0020), F(1,1247) = 1063.8, p < 0.0001, ω^2^ = 0.4597, respectively). Beta band power contained a significant amount of information during the sample (early and late sample, mean (± SEM): 0.0049 (± 0.0007), F(1,1247) = 200.02, p < 0.0001, ω^2^ = 0.1374 and 0.0065 (± 0.0008), F(1,1247) = 299.36, p < 0.0001, ω^2^ = 0.1928, respectively), and notable information during the delay (mid delay, mean (± SEM), 0.0121 (± 0.0011), F(1,1247) = 543.44, p < 0.0001, ω^2^ = 0.2993).

Other frequency bands (3-7 Hz ‘theta’, 8-12 Hz ‘alpha’, and 83-98 Hz ‘high gamma’) also contained information about the contralateral position during the sample phase and delay phases (SFig. 2). However, these frequency bands had much less information compared to the low gamma band (see Supplementary section 3). None of the frequency bands had meaningful information about the ipsilateral locations (refer to the extended data table 1 & 2 for a detailed overview). This location information contained in LFP power indicates involvement in processing the spatial component of the task, as binding each color to a location was necessary for localizing the change detection.

**Table 1:** Numerical values of position information.

| | average position information contralateral ($R_{adj.}^2$) | | | | | |
| --- | --- | --- | --- | --- | --- | --- |
| frequency | preSmp | earlySmp | lateSmp | earlyDly | midDly | lateDly |
| $\theta(3 - 7Hz)$ | 0.0001 | 0.0045 | 0.0096 | 0.0059 | 0.008 | 0.0046 |
| $\alpha(8 - 12Hz)$ | 0 | 0.0067 | 0.0088 | 0.0048 | 0.007 | 0.0037 |
| $\beta(13 - 19Hz)$ | 0.0008 | 0.0049 | 0.0065 | 0.0025 | 0.0121 | 0.0042 |
| $\gamma(33 - 48Hz)$ | 0.0004 | 0.0182 | 0.033 | 0.0073 | 0.0064 | 0.0017 |
| $\Gamma(83 - 98Hz)$ | -0.0003 | 0.0127 | 0.0152 | 0.0049 | 0.0028 | 0.0053 |
|  | standard error of the mean (contralateral) |  |  |  |  |  |
| $\theta(3 - 7Hz)$ | 0.0003 | 0.0006 | 0.001 | 0.0009 | 0.0009 | 0.0006 |
| $\alpha(8 - 12Hz)$ | 0.0003 | 0.0009 | 0.0011 | 0.0007 | 0.0008 | 0.0005 |
| $\beta(13 - 19Hz)$ | 0.0004 | 0.0007 | 0.0008 | 0.0005 | 0.0011 | 0.0005 |
| $\gamma(33 - 48Hz)$ | 0.0003 | 0.0019 | 0.002 | 0.0007 | 0.0007 | 0.0005 |
| $\Gamma(83 - 98Hz)$ | 0.0003 | 0.0021 | 0.0012 | 0.0007 | 0.0006 | 0.0007 |
| | effect size of test against null distribution ($\omega^2$) | | | | | |
| $\theta(3 - 7Hz)$ | -0.0006 | 0.1584 | 0.2323 | 0.1274 | 0.21 | 0.1384 |
| $\alpha(8 - 12Hz)$ | -0.0006 | 0.142 | 0.1811 | 0.1409 | 0.1899 | 0.1307 |
| $\beta(13 - 19Hz)$ | 0.0138 | 0.1374 | 0.1928 | 0.0691 | 0.2993 | 0.1716 |
| $\gamma(33 - 48Hz)$ | 0.0044 | 0.2327 | 0.4597 | 0.2584 | 0.2159 | 0.0419 |
| $\Gamma(83 - 98Hz)$ | 0.0014 | 0.1092 | 0.3477 | 0.1384 | 0.063 | 0.1576 |
| | average position information ipsilateral ($R_{adj.}^2$) | | | | | |
| $\theta(3 - 7Hz)$ | -0.0013 | 0.0013 | 0.0017 | 0.0058 | 0.0034 | 0.0018 |
| $\alpha(8 - 12Hz)$ | 0.0006 | 0.003 | 0.0013 | 0.0016 | 0.0005 | 0.0004 |
| $\beta(13 - 19Hz)$ | 0.0005 | 0.0017 | 0.0021 | 0.0004 | -0.0001 | 0.0004 |
| $\gamma(33 - 48Hz)$ | 0.0001 | 0.0006 | 0.0029 | 0.0025 | 0.0008 | 0.0007 |
| $\Gamma(83 - 98Hz)$ | -0.0001 | 0.0016 | 0.0009 | 0.0014 | 0.0007 | 0.0015 |
|  | standard error of the mean (ipsilateral) |  |  |  |  |  |
| $\theta(3 - 7Hz)$ | 0.0003 | 0.0005 | 0.0005 | 0.0007 | 0.0007 | 0.0004 |
| $\alpha(8 - 12Hz)$ | 0.0004 | 0.0005 | 0.0004 | 0.0005 | 0.0004 | 0.0004 |
| $\beta(13 - 19Hz)$ | 0.0003 | 0.0004 | 0.0004 | 0.0004 | 0.0003 | 0.0004 |
| $\gamma(33 - 48Hz)$ | 0.0004 | 0.0004 | 0.0004 | 0.0005 | 0.0004 | 0.0003 |
| $\Gamma(83 - 98Hz)$ | 0.0003 | 0.0005 | 0.0004 | 0.0005 | 0.0004 | 0.0004 |
| | effect size of test against null distribution ($\omega^2$) | | | | | |
| $\theta(3 - 7Hz)$ | 0.0689 | 0.0218 | 0.0399 | 0.196 | 0.0736 | 0.0566 |
| $\alpha(8 - 12Hz)$ | 0.0072 | 0.1012 | 0.0256 | 0.0352 | 0.0055 | 0.0042 |
| $\beta(13 - 19Hz)$ | 0.0092 | 0.0531 | 0.0695 | 0.0033 | -0.0005 | 0.0033 |
| $\gamma(33 - 48Hz)$ | -0.0007 | 0.0051 | 0.1166 | 0.0801 | 0.0123 | 0.0125 |
| $\Gamma(83 - 98Hz)$ | -0.0005 | 0.0376 | 0.0172 | 0.025 | 0.0097 | 0.038 |

**Table 2:** Statistics of position information.

| ID | F | $df_1$ | $df_2$ | p |
| --- | --- | --- | --- | --- |
| $\theta(3 - 7Hz)contraPreSmp$ | 0.3103 | 1 | 1247 | 0.5776 |
| $\theta(3 - 7Hz)contraEarlySmp$ | 236.1105 | 1 | 1247 | < 0.0001 |
| $\theta(3 - 7Hz)contraLateSmp$ | 378.9953 | 1 | 1247 | < 0.0001 |
| $\theta(3 - 7Hz)contraEarlyDly$ | 183.325 | 1 | 1247 | < 0.0001 |
| $\theta(3 - 7Hz)contraMidDly$ | 332.9131 | 1 | 1247 | < 0.0001 |
| $\theta(3 - 7Hz)contraLateDlyEarlyChc$ | 201.582 | 1 | 1247 | < 0.0001 |
| $\theta(3 - 7Hz)ipsiPreSmp$ | 93.4328 | 1 | 1247 | < 0.0001 |
| $\theta(3 - 7Hz)ipsiEarlySmp$ | 28.8243 | 1 | 1247 | < 0.0001 |
| $\theta(3 - 7Hz)ipsiLateSmp$ | 52.9692 | 1 | 1247 | < 0.0001 |
| $\theta(3 - 7Hz)ipsiEarlyDly$ | 305.5275 | 1 | 1247 | < 0.0001 |
| $\theta(3 - 7Hz)ipsiMidDly$ | 100.2107 | 1 | 1247 | < 0.0001 |
| $\theta(3 - 7Hz)ipsiLateDlyEarlyChc$ | 75.9505 | 1 | 1247 | < 0.0001 |
| $\alpha(8 - 12Hz)contraPreSmp$ | 0.196 | 1 | 1247 | 0.6581 |
| $\alpha(8 - 12Hz)contraEarlySmp$ | 207.6518 | 1 | 1247 | < 0.0001 |
| $\alpha(8 - 12Hz)contraLateSmp$ | 277.1249 | 1 | 1247 | < 0.0001 |
| $\alpha(8 - 12Hz)contraEarlyDly$ | 205.7884 | 1 | 1247 | < 0.0001 |
| $\alpha(8 - 12Hz)contraMidDly$ | 293.8236 | 1 | 1247 | < 0.0001 |
| $\alpha(8 - 12Hz)contraLateDlyEarlyChc$ | 188.7494 | 1 | 1247 | < 0.0001 |
| $\alpha(8 - 12Hz)ipsiPreSmp$ | 10.015 | 1 | 1247 | 0.0016 |
| $\alpha(8 - 12Hz)ipsiEarlySmp$ | 141.6304 | 1 | 1247 | < 0.0001 |
| $\alpha(8 - 12Hz)ipsiLateSmp$ | 33.8392 | 1 | 1247 | < 0.0001 |
| $\alpha(8 - 12Hz)ipsiEarlyDly$ | 46.5335 | 1 | 1247 | < 0.0001 |
| $\alpha(8 - 12Hz)ipsiMidDly$ | 7.9221 | 1 | 1247 | 0.005 |
| $\alpha(8 - 12Hz)ipsiLateDlyEarlyChc$ | 6.2923 | 1 | 1247 | 0.0123 |
| $\beta(13 - 19Hz)contraPreSmp$ | 18.4705 | 1 | 1247 | < 0.0001 |
| $\beta(13 - 19Hz)contraEarlySmp$ | 200.0195 | 1 | 1247 | < 0.0001 |
| $\beta(13 - 19Hz)contraLateSmp$ | 299.3604 | 1 | 1247 | < 0.0001 |
| $\beta(13 - 19Hz)contraEarlyDly$ | 93.7528 | 1 | 1247 | < 0.0001 |
| $\beta(13 - 19Hz)contraMidDly$ | 534.4364 | 1 | 1247 | < 0.0001 |
| $\beta(13 - 19Hz)contraLateDlyEarlyChc$ | 259.699 | 1 | 1247 | < 0.0001 |
| $\beta(13 - 19Hz)ipsiPreSmp$ | 12.6119 | 1 | 1247 | 0.0004 |
| $\beta(13 - 19Hz)ipsiEarlySmp$ | 71.0949 | 1 | 1247 | < 0.0001 |
| $\beta(13 - 19Hz)ipsiLateSmp$ | 94.2461 | 1 | 1247 | < 0.0001 |
| $\beta(13 - 19Hz)ipsiEarlyDly$ | 5.0947 | 1 | 1247 | 0.0242 |
| $\beta(13 - 19Hz)ipsiMidDly$ | 0.4002 | 1 | 1247 | 0.5271 |
| $\beta(13 - 19Hz)ipsiLateDlyEarlyChc$ | 5.1808 | 1 | 1247 | 0.023 |
| $\gamma(33 - 48Hz)contraPreSmp$ | 6.4733 | 1 | 1247 | 0.0111 |
| $\gamma(33 - 48Hz)contraEarlySmp$ | 379.8323 | 1 | 1247 | < 0.0001 |
| $\gamma(33 - 48Hz)contraLateSmp$ | 1063.827 | 1 | 1247 | < 0.0001 |
| $\gamma(33 - 48Hz)contraEarlyDly$ | 436.2718 | 1 | 1247 | < 0.0001 |
| $\gamma(33 - 48Hz)contraMidDly$ | 344.8669 | 1 | 1247 | < 0.0001 |
| $\gamma(33 - 48Hz)contraLateDlyEarlyChc$ | 55.5925 | 1 | 1247 | < 0.0001 |
| $\gamma(33 - 48Hz)ipsiPreSmp$ | 0.1014 | 1 | 1247 | 0.7502 |
| $\gamma(33 - 48Hz)ipsiEarlySmp$ | 7.44 | 1 | 1247 | 0.0065 |
| $\gamma(33 - 48Hz)ipsiLateSmp$ | 165.8605 | 1 | 1247 | < 0.0001 |
| $\gamma(33 - 48Hz)ipsiEarlyDly$ | 109.7774 | 1 | 1247 | < 0.0001 |
| $\gamma(33 - 48Hz)ipsiMidDly$ | 16.5156 | 1 | 1247 | < 0.0001 |
| $\gamma(33 - 48Hz)ipsiLateDlyEarlyChc$ | 16.8524 | 1 | 1247 | < 0.0001 |
| Table 2: continued |  |  |  |  |
| $\Gamma(83 - 98Hz)_{contraPreSmp}$ | 2.7801 | 1 | 1247 | 0.0957 |
| $\Gamma(83 - 98Hz)_{contraEarlySmp}$ | 154.0675 | 1 | 1247 | < 0.0001 |
| $\Gamma(83 - 98Hz)_{contraLateSmp}$ | 666.6498 | 1 | 1247 | < 0.0001 |
| $\Gamma(83 - 98Hz)_{contraEarlyDly}$ | 201.7003 | 1 | 1247 | < 0.0001 |
| $\Gamma(83 - 98Hz)_{contraMidDly}$ | 85.0001 | 1 | 1247 | < 0.0001 |
| $\Gamma(83 - 98Hz)_{contraLateDlyEarlyChc}$ | 234.6346 | 1 | 1247 | < 0.0001 |
| $\Gamma(83 - 98Hz)_{ipsiPreSmp}$ | 0.3795 | 1 | 1247 | 0.538 |
| $\Gamma(83 - 98Hz)_{ipsiEarlySmp}$ | 49.7564 | 1 | 1247 | < 0.0001 |
| $\Gamma(83 - 98Hz)_{ipsiLateSmp}$ | 22.8253 | 1 | 1247 | < 0.0001 |
| $\Gamma(83 - 98Hz)_{ipsiEarlyDly}$ | 32.9647 | 1 | 1247 | < 0.0001 |
| $\Gamma(83 - 98Hz)_{ipsiMidDly}$ | 13.1832 | 1 | 1247 | 0.0003 |
| $\Gamma(83 - 98Hz)_{ipsiLateDlyEarlyChc}$ | 50.3882 | 1 | 1247 | < 0.0001 |

### Working memory load modulated gamma

The major manipulation affecting cognitive processing in our task was the number of squares the birds had to memorize as it determined the load of WM. We considered three load conditions (‘loads’). Because power contained information only for contralateral locations, we analyzed load effects for the number of squares presented in the visual hemifield contralateral to the recording electrode. Trials in which only one square was presented during the sample (Fig. 1A), were considered to have ‘load 1’ (irrespective of the number of squares on the other side of the screen). Following this logic, trials with two, or three presented colors were considered ‘load 2’, and ‘load 3’, respectively. To understand how LFP-power was modulated by WM-load, we again compared the power of all gamma-modulated electrodes during the sample and memory delay phases to baseline power during the inter-trial interval. When comparing power across the different loads, the local maximum of power in the low gamma band appeared to be modulated, with higher loads reducing average power (Fig. 3E, SFig. 3). Similarly, the power in the lower bands appeared to be affected by load. To better quantify the load effect, we tested its effect on power in the five major frequency bands introduced above (we focus on the low gamma and the beta band that prominently affected by the overall task, refer to SFig. 4 for the other frequency bands). The mean power in the respective frequency band, across all channels with significant gamma power modulation in load 1 trials, was compared by calculating the average change in power per added item. Power in the low gamma band decreased as load increased during sample and delay but reversed this modulation during the subsequent choice phase (Fig. 3C). The beta band showed the opposite effect of load, with power generally increasing at higher loads. We further quantified the magnitude of the load effect by calculating the effect size (PEV, ω^2^) of the LFP differences for different loads over time (see methods for details). The influence of load on low-gamma power was largest towards the end of the sample (power decreased with load), and in the choice phase (power increased with load). The strongest beta power load modulation started appearing during the middle of the delay phase (power increased with load), peaking at the end of the delay (Fig. 3D, refer to table 3 for numerical values). This means that LFP power was substantially affected by both the locations of the presented stimuli and by the WM load. Therefore, LFP processes seem to be tightly linked to ongoing cognitive processing of the WM task, during both sample encoding of memory items, and their subsequent maintenance during the delay.

**Table 3:** Avg. chg per item, numerical values and statistics

| bin start relative to sample on (ms) | avg. chg. per item ($\times 10^{-2}\%$) ±SEM, test vs. 0; effect size of factor load |
| --- | --- |
| $\theta(3 - 7Hz)$ | |
| -200 | -0.0002 ± 0.028, t(288) = 0.01833, p = 0.9854; $\omega^2$ = 0.0059 |
| -100 | 0.0016 ± 0.0235, t(288) = -0.18974, p = 0.8496; $\omega^2$ = 0.0032 |
| 0 | 0.0055 ± 0.0202, t(288) = -0.74963, p = 0.4541; $\omega^2$ = -0.0017 |
| 100 | 0.008 ± 0.017, t(288) = -1.2945, p = 0.1965; $\omega^2$ = -0.0026 |
| 200 | 0.0095 ± 0.014, t(288) = -1.8655, p = 0.0631; $\omega^2$ = 0.002 |
| 300 | 0.0103 ± 0.0122, t(288) = -2.3301, p = 0.0205; $\omega^2$ = 0.0019 |
| 400 | 0.0124 ± 0.0129, t(288) = -2.6689, p = 0.008; $\omega^2$ = -0.0262 |
| 500 | 0.0198 ± 0.0155, t(288) = -3.5227, p = 0.0005; $\omega^2$ = -0.0374 |
| 600 | 0.0318 ± 0.0174, t(288) = -5.039, p < 0.0001; $\omega^2$ = -0.0126 |
| 700 | 0.0429 ± 0.0169, t(288) = -6.9947, p < 0.0001; $\omega^2$ = 0.0209 |
| 800 | 0.0486 ± 0.015, t(288) = -8.9559, p < 0.0001; $\omega^2$ = 0.0412 |
| 900 | 0.0492 ± 0.0131, t(288) = -10.3168, p < 0.0001; $\omega^2$ = 0.045 |
| 1000 | 0.0487 ± 0.0122, t(288) = -11.0283, p < 0.0001; $\omega^2$ = 0.0465 |
| 1100 | 0.0517 ± 0.0122, t(288) = -11.7262, p < 0.0001; $\omega^2$ = 0.0584 |
| 1200 | 0.0592 ± 0.0129, t(288) = -12.6267, p < 0.0001; $\omega^2$ = 0.0865 |
| 1300 | 0.0718 ± 0.0144, t(288) = -13.7098, p < 0.0001; $\omega^2$ = 0.1366 |
| 1400 | 0.0918 ± 0.0168, t(288) = -15.0213, p < 0.0001; $\omega^2$ = 0.2025 |
| 1500 | 0.1182 ± 0.0197, t(288) = -16.537, p < 0.0001; $\omega^2$ = 0.2471 |
| 1600 | 0.1423 ± 0.0226, t(288) = -17.3278, p < 0.0001; $\omega^2$ = 0.2502 |
| 1700 | 0.1503 ± 0.0248, t(288) = -16.7352, p < 0.0001; $\omega^2$ = 0.2248 |
| 1800 | 0.1339 ± 0.0245, t(288) = -15.0652, p < 0.0001; $\omega^2$ = 0.1833 |
| 1900 | 0.1003 ± 0.0242, t(288) = -11.4106, p < 0.0001; $\omega^2$ = 0.1308 |
| 2000 | 0.0714 ± 0.0297, t(288) = -6.6298, p < 0.0001; $\omega^2$ = 0.0859 |
| 2100 | 0.0537 ± 0.0334, t(287) = -4.4192, p < 0.0001; $\omega^2$ = 0.0529 |
| $\alpha(8 - 12Hz)$ | |
| -200 | -0.0004 ± 0.0257, t(268) = 0.038799, p = 0.9691; $\omega^2$ = 0.006 |
| -100 | 0.01 ± 0.0215, t(268) = -1.2311, p = 0.2194; $\omega^2$ = 0.0003 |
| 0 | 0.0217 ± 0.0187, t(269) = -3.0882, p = 0.0022; $\omega^2$ = 0.0034 |
| 100 | 0.0276 ± 0.015, t(269) = -4.9084, p < 0.0001; $\omega^2$ = 0.0289 |
| 200 | 0.0352 ± 0.0133, t(269) = -7.0544, p < 0.0001; $\omega^2$ = 0.0197 |
| 300 | 0.0227 ± 0.0147, t(269) = -4.1294, p < 0.0001; $\omega^2$ = 0.0056 |
| 400 | 0.0112 ± 0.0209, t(269) = -1.4248, p = 0.1554; $\omega^2$ = -0.0133 |
| 500 | 0.0154 ± 0.0264, t(269) = -1.5565, p = 0.1208; $\omega^2$ = -0.0437 |
| 600 | 0.034 ± 0.0305, t(269) = -2.9688, p = 0.0033; $\omega^2$ = -0.0223 |
| 700 | 0.0521 ± 0.0292, t(269) = -4.7633, p < 0.0001; $\omega^2$ = 0.0062 |
| 800 | 0.065 ± 0.021, t(269) = -8.2519, p < 0.0001; $\omega^2$ = 0.0199 |
| 900 | 0.0681 ± 0.016, t(269) = -11.324, p < 0.0001; $\omega^2$ = 0.048 |
| 1000 | 0.0461 ± 0.014, t(269) = -8.7594, p < 0.0001; $\omega^2$ = 0.0126 |
| 1100 | 0.0384 ± 0.0147, t(269) = -6.98, p < 0.0001; $\omega^2$ = 0.0016 |
| 1200 | 0.0448 ± 0.0146, t(269) = -8.1937, p < 0.0001; $\omega^2$ = -0.002 |
| 1300 | 0.0631 ± 0.0154, t(269) = -10.8933, p < 0.0001; $\omega^2$ = -0.0074 |
| 1400 | 0.1071 ± 0.0184, t(269) = -15.5377, p < 0.0001; $\omega^2$ = 0.066 |
| 1500 | 0.1502 ± 0.0205, t(269) = -19.5323, p < 0.0001; $\omega^2$ = 0.1728 |
| 1600 | 0.1909 ± 0.0257, t(269) = -19.8154, p < 0.0001; $\omega^2$ = 0.1969 |
| 1700 | 0.2037 ± 0.0268, t(269) = -20.2351, p < 0.0001; $\omega^2$ = 0.2055 |
| 1800 | 0.1787 ± 0.0246, t(269) = -19.3425, p < 0.0001; $\omega^2$ = 0.1967 |
| 1900 | 0.143 ± 0.0213, t(269) = -17.8667, p < 0.0001; $\omega^2$ = 0.1 |
| 2000 | 0.1168 ± 0.0221, t(269) = -14.0688, p < 0.0001; $\omega^2$ = 0.0719 |
| 2100 | 0.0953 ± 0.0201, t(269) = -12.643, p < 0.0001; $\omega^2$ = 0.0656 |

| Table 3: continued |  |
| --- | --- |
| $\beta(13 - 19Hz)$ | |
| -200 | $0.0067 \pm 0.0391$, $t(191) = -0.38236$, $p = 0.7026$; $\omega^2 = 0.0131$ |
| -100 | $-0.0061 \pm 0.0221$, $t(192) = 0.62171$, $p = 0.5349$; $\omega^2 = 0.0147$ |
| 0 | $0.0122 \pm 0.0201$, $t(192) = -1.3742$, $p = 0.171$; $\omega^2 = -0.0003$ |
| 100 | $0.0323 \pm 0.0132$, $t(192) = -5.5143$, $p < 0.0001$; $\omega^2 = 0.011$ |
| 200 | $0.0125 \pm 0.0114$, $t(192) = -2.4815$, $p = 0.0139$; $\omega^2 = 0.0265$ |
| 300 | $-0.0354 \pm 0.0132$, $t(192) = 6.0399$, $p < 0.0001$; $\omega^2 = 0.058$ |
| 400 | $-0.0225 \pm 0.0161$, $t(192) = 3.1334$, $p = 0.002$; $\omega^2 = 0.0217$ |
| 500 | $-0.0262 \pm 0.018$, $t(192) = 3.2814$, $p = 0.0012$; $\omega^2 = 0.0202$ |
| 600 | $-0.0175 \pm 0.0177$, $t(192) = 2.2277$, $p = 0.0271$; $\omega^2 = 0.0164$ |
| 700 | $-0.0089 \pm 0.0167$, $t(192) = 1.2002$, $p = 0.2315$; $\omega^2 = 0.0154$ |
| 800 | $0.0059 \pm 0.0145$, $t(192) = -0.91096$, $p = 0.3635$; $\omega^2 = 0.0232$ |
| 900 | $0.0236 \pm 0.0133$, $t(192) = -4.0074$, $p < 0.0001$; $\omega^2 = 0.0184$ |
| 1000 | $0.0002 \pm 0.0117$, $t(192) = -0.033248$, $p = 0.9735$; $\omega^2 = 0.0044$ |
| 1100 | $0.0061 \pm 0.0123$, $t(192) = -1.1262$, $p = 0.2615$; $\omega^2 = 0.0042$ |
| 1200 | $0.0212 \pm 0.0113$, $t(192) = -4.2481$, $p < 0.0001$; $\omega^2 = 0.0007$ |
| 1300 | $0.0403 \pm 0.0118$, $t(192) = -7.7319$, $p < 0.0001$; $\omega^2 = 0.0044$ |
| 1400 | $0.0721 \pm 0.0154$, $t(192) = -10.5736$, $p < 0.0001$; $\omega^2 = 0.0368$ |
| 1500 | $0.0977 \pm 0.0156$, $t(192) = -14.1347$, $p < 0.0001$; $\omega^2 = 0.0725$ |
| 1600 | $0.1319 \pm 0.0162$, $t(192) = -18.3303$, $p < 0.0001$; $\omega^2 = 0.118$ |
| 1700 | $0.1321 \pm 0.0181$, $t(192) = -16.4302$, $p < 0.0001$; $\omega^2 = 0.0797$ |
| 1800 | $0.1126 \pm 0.0154$, $t(192) = -16.4416$, $p < 0.0001$; $\omega^2 = 0.0787$ |
| 1900 | $0.0988 \pm 0.0143$, $t(192) = -15.5418$, $p < 0.0001$; $\omega^2 = 0.0513$ |
| 2000 | $0.1023 \pm 0.0147$, $t(192) = -15.7297$, $p < 0.0001$; $\omega^2 = 0.085$ |
| 2100 | $0.0714 \pm 0.0176$, $t(192) = -9.1154$, $p < 0.0001$; $\omega^2 = 0.0485$ |
| $\gamma(33 - 48Hz)$ | |
| -200 | $0.0284 \pm 0.0424$, $t(248) = -1.7162$, $p = 0.0874$; $\omega^2 = 0.0084$ |
| -100 | $-0.0237 \pm 0.0183$, $t(248) = 3.3185$, $p = 0.001$; $\omega^2 = 0.016$ |
| 0 | $-0.0176 \pm 0.0374$, $t(248) = 1.2047$, $p = 0.2295$; $\omega^2 = 0.0062$ |
| 100 | $0.0117 \pm 0.0086$, $t(248) = -3.4988$, $p = 0.0006$; $\omega^2 = 0.0313$ |
| 200 | $-0.0451 \pm 0.0082$, $t(248) = 14.0087$, $p < 0.0001$; $\omega^2 = 0.0592$ |
| 300 | $-0.0514 \pm 0.0107$, $t(248) = 12.297$, $p < 0.0001$; $\omega^2 = 0.0485$ |
| 400 | $-0.0414 \pm 0.0138$, $t(248) = 7.6807$, $p < 0.0001$; $\omega^2 = 0.0244$ |
| 500 | $-0.0527 \pm 0.0139$, $t(248) = 9.7$, $p < 0.0001$; $\omega^2 = 0.0453$ |
| 600 | $-0.0471 \pm 0.0095$, $t(248) = 12.7011$, $p < 0.0001$; $\omega^2 = 0.0838$ |
| 700 | $-0.0497 \pm 0.0083$, $t(248) = 15.2964$, $p < 0.0001$; $\omega^2 = 0.0716$ |
| 800 | $-0.0306 \pm 0.0076$, $t(248) = 10.2899$, $p < 0.0001$; $\omega^2 = 0.0584$ |
| 900 | $-0.0015 \pm 0.0076$, $t(248) = 0.51818$, $p = 0.6048$; $\omega^2 = 0.0098$ |
| 1000 | $-0.0053 \pm 0.0084$, $t(248) = 1.6083$, $p = 0.109$; $\omega^2 = 0.0387$ |
| 1100 | $-0.0153 \pm 0.0086$, $t(248) = 4.578$, $p < 0.0001$; $\omega^2 = 0.0262$ |
| 1200 | $-0.0184 \pm 0.0074$, $t(248) = 6.3718$, $p < 0.0001$; $\omega^2 = 0.0044$ |
| 1300 | $-0.0282 \pm 0.0092$, $t(248) = 7.8115$, $p < 0.0001$; $\omega^2 = 0.0172$ |
| 1400 | $-0.0379 \pm 0.0102$, $t(248) = 9.54$, $p < 0.0001$; $\omega^2 = -0.0012$ |
| 1500 | $-0.027 \pm 0.01$, $t(248) = 6.9461$, $p < 0.0001$; $\omega^2 = 0.0096$ |
| 1600 | $-0.0199 \pm 0.0099$, $t(248) = 5.1705$, $p < 0.0001$; $\omega^2 = 0.0152$ |
| 1700 | $0.0107 \pm 0.0101$, $t(248) = -2.6985$, $p = 0.0074$; $\omega^2 = 0.0446$ |
| 1800 | $0.0187 \pm 0.0107$, $t(248) = -4.452$, $p < 0.0001$; $\omega^2 = 0.0499$ |
| 1900 | $0.0722 \pm 0.0101$, $t(248) = -18.2718$, $p < 0.0001$; $\omega^2 = 0.0711$ |
| 2000 | $0.089 \pm 0.0108$, $t(248) = -21.1006$, $p < 0.0001$; $\omega^2 = 0.0799$ |
| 2100 | $0.0786 \pm 0.0106$, $t(248) = -18.974$, $p < 0.0001$; $\omega^2 = 0.0525$ |
| $\Gamma(83 - 98Hz)$ | |
| -200 | $0.0176 \pm 0.0364$, $t(229) = -1.1913$, $p = 0.2348$; $\omega^2 = 0.0056$ |
| -100 | $-0.0091 \pm 0.0275$, $t(233) = 0.82159$, $p = 0.4121$; $\omega^2 = 0.0148$ |
| 0 | $-0.0183 \pm 0.0211$, $t(230) = 2.1443$, $p = 0.0331$; $\omega^2 = 0.0103$ |
| 100 | $-0.0491 \pm 0.0057$, $t(233) = 21.2992$, $p < 0.0001$; $\omega^2 = 0.0797$ |
| 200 | $-0.0532 \pm 0.0067$, $t(233) = 19.596$, $p < 0.0001$; $\omega^2 = 0.0842$ |
| 300 | $-0.002 \pm 0.0071$, $t(233) = 0.68699$, $p = 0.4928$; $\omega^2 = 0.0389$ |
| 400 | $-0.0159 \pm 0.0073$, $t(233) = 5.3819$, $p < 0.0001$; $\omega^2 = 0.029$ |
| 500 | $-0.0288 \pm 0.008$, $t(233) = 8.8849$, $p < 0.0001$; $\omega^2 = 0.0434$ |
| 600 | $-0.0059 \pm 0.0096$, $t(233) = 1.536$, $p = 0.1259$; $\omega^2 = 0.0284$ |
| 700 | $-0.0118 \pm 0.0076$, $t(233) = 3.8498$, $p = 0.0002$; $\omega^2 = 0.0319$ |
| 800 | $-0.0049 \pm 0.0083$, $t(233) = 1.4611$, $p = 0.1453$; $\omega^2 = 0.0272$ |
| 900 | $-0.0127 \pm 0.0066$, $t(233) = 4.8156$, $p < 0.0001$; $\omega^2 = 0.0282$ |
| 1000 | $0.0011 \pm 0.0088$, $t(233) = -0.30018$, $p = 0.7643$; $\omega^2 = 0.0133$ |
| 1100 | $0.0005 \pm 0.0081$, $t(233) = -0.14972$, $p = 0.8811$; $\omega^2 = 0.0093$ |
| 1200 | $-0.0111 \pm 0.0063$, $t(233) = 4.3941$, $p < 0.0001$; $\omega^2 = 0.0099$ |
| 1300 | $-0.0048 \pm 0.0057$, $t(233) = 2.0878$, $p = 0.0379$; $\omega^2 = 0.0051$ |
| 1400 | $-0.0146 \pm 0.007$, $t(233) = 5.1816$, $p < 0.0001$; $\omega^2 = 0.0011$ |
| 1500 | $-0.0259 \pm 0.0071$, $t(233) = 9.0215$, $p < 0.0001$; $\omega^2 = -0.0062$ |
| 1600 | $-0.0367 \pm 0.0077$, $t(233) = 11.8365$, $p < 0.0001$; $\omega^2 = -0.0103$ |
| 1700 | $-0.0302 \pm 0.007$, $t(233) = 10.7248$, $p < 0.0001$; $\omega^2 = -0.0225$ |
| 1800 | $-0.0325 \pm 0.0068$, $t(233) = 11.9615$, $p < 0.0001$; $\omega^2 = 0.0216$ |
| 1900 | $-0.0206 \pm 0.006$, $t(233) = 8.4922$, $p < 0.0001$; $\omega^2 = 0.0336$ |
| 2000 | $-0.0418 \pm 0.0097$, $t(233) = 10.6327$, $p < 0.0001$; $\omega^2 = 0.0104$ |
| 2100 | $0.002 \pm 0.0072$, $t(233) = -0.69613$, $p = 0.487$; $\omega^2 = 0.0109$ |

**Figure 4:**
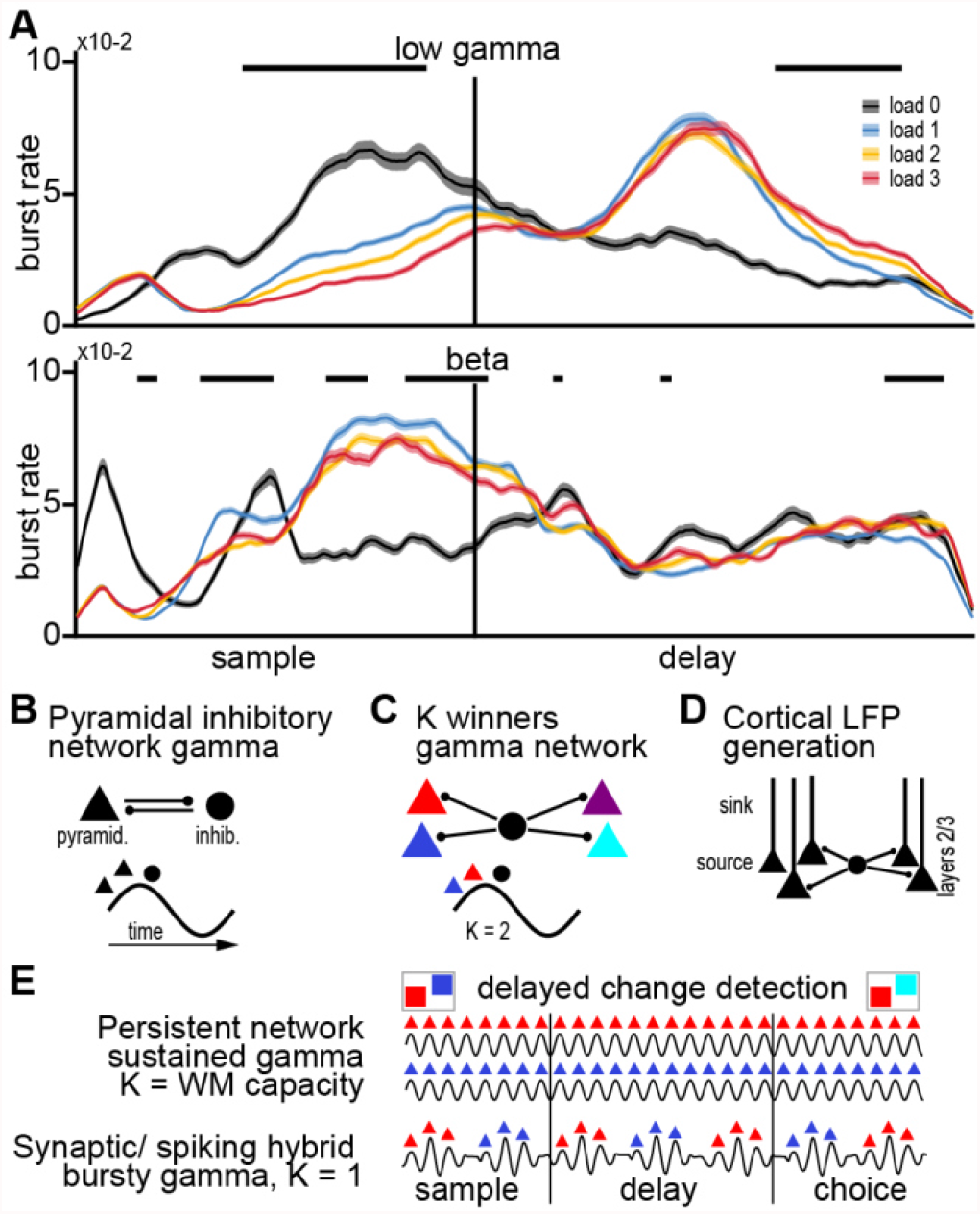
(**A**) Trial burst rate of low gamma and beta frequency bands during the trial at gamma modulated sites. Burst rate of low gamma strongly increases towards the end of the sample phase, while beta has peak burst rate in the middle of the delay. Load modulation occurs with higher loads decreasing burst rate in the sample but increasing burst rate towards the end of the delay. Black bars indicate consecutive significance between loads 1-3 (p < 0.05) over 2 cycles of the bands center frequency (**B**) Schematic generation of mammalian pyramidal inhibitory network gamma (PING). Gamma oscillation is generated in a cycle when excitatory pyramidal cells first become active, exciting inhibitory parvalbumin positive interneurons that provide dense, short-lasting feedback inhibition. The inhibition briefly shuts down the pyramidal cells to terminate the cycle. (**C**) Implementation of a winner-take-all dynamic. If several pyramidal populations (colored triangles) are connected to the same inhibitory population (black circle), the gamma generating feedback inhibition can implement a K winners take all dynamic where only the K most excited populations will spike before the feedback inhibition deactivates all populations. For example, the earlier spiking of blue and red in each cycle results in K = 2. (**D**) Cortical layer organization facilitate gamma oscillation. Many similarly aligned pyramidal cells receive rhythmic, peri-somatic inhibition. The pyramidal cells are thought to act as aligned dipoles with the source close to the somas and the sink in the apical dendrites, creating an extracellular field. The gamma in cortical LFPs is thus generated in the superficial layers of cortex. Crow NCL lacks this layered anatomical organization. (**E**) Two different networks solving a 2-item delay change detection task. The two colored squares can be retained either by selective, persistent activity (top) in a network where gamma implements a K = WM capacity winner take all algorithm, or alternatively, in a network relying both on intermittent spiking and synaptic mechanisms with K = 1. In the latter, since K = 1, the two memory representations take turn being active and silent resulting in bursting gamma. In the silent periods, information is retained in synaptic changes rather than sustained spiking.

### Beta and Gamma appear in bursts

An additional observation we made was that power modulations in the low gamma band appeared as bursts throughout sample and delay phase (Fig. 1B). In a study in which monkeys performed a sequential version of our task (Lundqvist et al., 2016), increases in gamma power were found to originate from sparse and temporarily defined ‘bursts’ of power. We tested if the increase in gamma power was due to individual bursts by investigating the potential burst events (power crossing a threshold of mean+1.5*SD for two cycles). We calculated burst rates over time (i.e., the observed rate of bursts at any given time in a trial, see methods for details). Burst rate in the low gamma band increased throughout the sample phase, peaking in the late sample phase (load 1, mean ± SEM, 0.0834 ± 0.0008 at 620 ms), before gradually reducing throughout the delay (Fig. 4A top, table 4, SFig. 4 for alpha and high gamma). Notably, the burst rate increased again during the latest part of the delay. These burst rates were also load-dependent in two directions depending on the task phase. During the sample phase, the burst rate significantly decreased with load, while during the late delay phase the burst rate increased with load (Fig. 4A, refer to table 5 for statistical values). The beta band also showed this effect (refer to Supplementary section 4 for the other frequency bands and for a comparison of bursts to population spiking rate). The phase and load-dependent rates of low gamma and beta bursts correlated with processing demands of WM for encoding during the sample, maintenance during the delay, and preparation for decoding towards the end of the delay and choice phase.

**Table 4:**
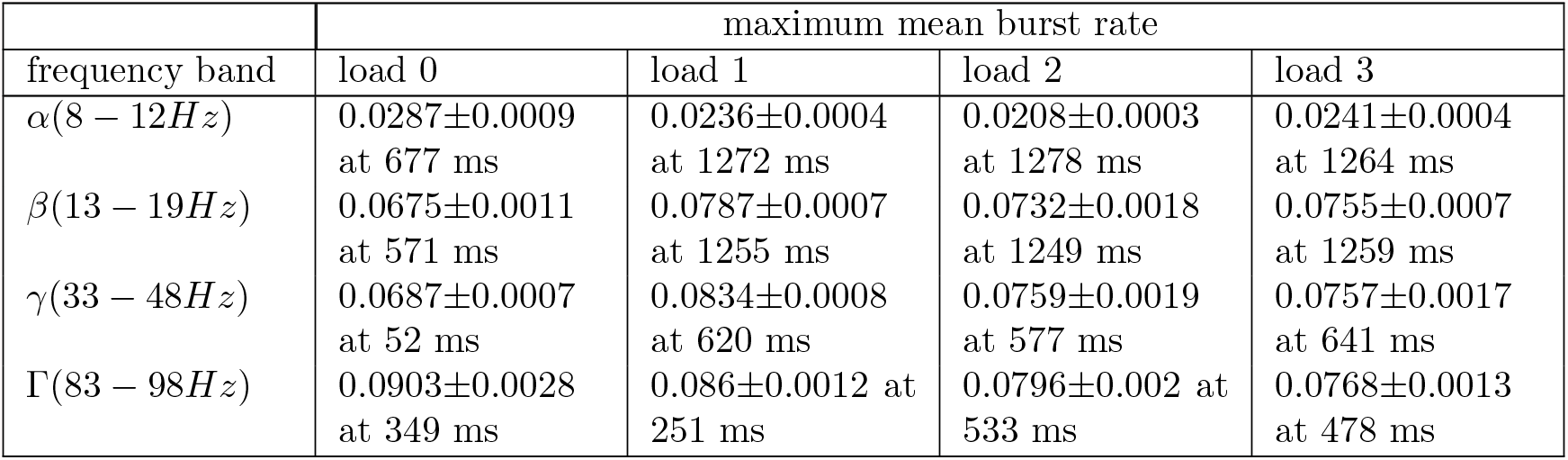
Numerical values of maxima of burst rates.

| frequency band | maximum mean burst rate |  |  |  |
| --- | --- | --- | --- | --- |
|  | load 0 | load 1 | load 2 | load 3 |
| $\alpha(8 - 12Hz)$ | $0.0287 \pm 0.0009$<br>at 677 ms | $0.0236 \pm 0.0004$<br>at 1272 ms | $0.0208 \pm 0.0003$<br>at 1278 ms | $0.0241 \pm 0.0004$<br>at 1264 ms |
| $\beta(13 - 19Hz)$ | $0.0675 \pm 0.0011$<br>at 571 ms | $0.0787 \pm 0.0007$<br>at 1255 ms | $0.0732 \pm 0.0018$<br>at 1249 ms | $0.0755 \pm 0.0007$<br>at 1259 ms |
| $\gamma(33 - 48Hz)$ | $0.0687 \pm 0.0007$<br>at 52 ms | $0.0834 \pm 0.0008$<br>at 620 ms | $0.0759 \pm 0.0019$<br>at 577 ms | $0.0757 \pm 0.0017$<br>at 641 ms |
| $\Gamma(83 - 98Hz)$ | $0.0903 \pm 0.0028$<br>at 349 ms | $0.086 \pm 0.0012$ at<br>251 ms | $0.0796 \pm 0.002$ at<br>533 ms | $0.0768 \pm 0.0013$<br>at 478 ms |

**Table 5:**
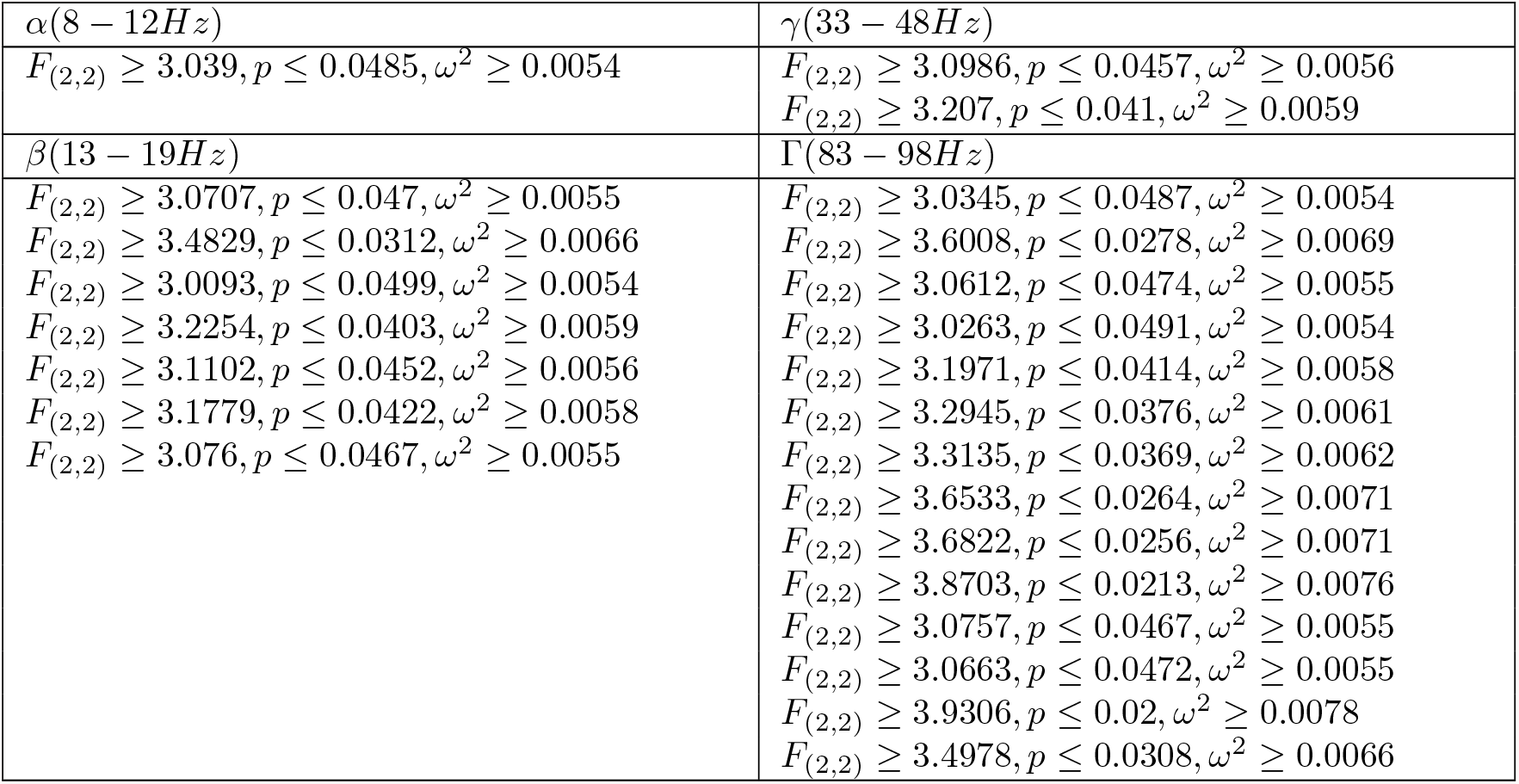
Statistics of burst rates. Values of the 1-way ANOVA for load. Each row reports the relevant values of the significant phases indicated in Fig. 4A and SFig. 4., ordered chronologically.

## Discussion

We observed cognitively modulated oscillations of LFP in the NCL of carrion crows performing a WM task. Oscillations occurred in a narrow gamma band, and in the beta band. This data shares many similarities with those observed in monkey PFC. While these results are consistent with behavioral and single-cell observations they are remarkable given that WM of birds and monkeys have diverging neuronal architectures that evolved independently over the last 320 million years (Benton & Donoghue, 2007).

### Cognitively modulated non-cortical gamma

The laminar and columnar organization of the mammalian cortex, with similarly aligned pyramidal cells, is thought to produce extracellular electrical fields that facilitate the observation of rhythmic population activity (Fig. 4B; (Buzsáki et al., 2012; Einevoll et al., 2013)). However, the associative pallium of birds lacks this structure entirely (Güntürkün & Bugnyar, 2016), and the mosaic-like arrangement of fiber patches in NCL (Stacho et al., 2020) differs substantially from the highly structured, layered, organization of the PFC. Gamma oscillations in birds have first been reported in the optic tectum of pigeons (Neuenschwander & Varela, 1993) and barn owls (Sridharan et al., 2011), a midbrain structure that like the mammalian neocortex displays a separation between grey and white matter and organization into highly structured layers (Güntürkün et al., 2020). Only recently modulated gamma was reported in the (non-layered) telencephalon of birds. In the song system of singing zebra finches (Brown et al., 2021; Lewandowski & Schmidt, 2011), and in the hippocampus of sleeping zebra finches (van der Meij et al., 2020). Functionally involved in such gamma oscillations are excitatory cell types, homologous to mammalian excitatory neurons, which are part of neuronal circuitry that can be optogenetically induced to produce broad range gamma oscillations (Spool et al., 2021), Fig. 4B). These neurons were found in the zebra finch pallium, adjacent to NCL (Spool et al., 2021). Based on the similarities of the observations of gamma in the avian optic tectum it has been suggested that gamma rhythms play an essential role in information processing and are thus evolutionary conserved (Sridharan & Knudsen, 2015). We now demonstrate that the NCL of crows also shows gamma modulation of the LFP, importantly in the absence of motor planning or execution, and directly linked to cognition. This is despite the anatomical differences between the layered PFC and nuclear NCL, in terms of the architecture of the telencephalon at this mesoscale. Therefore, the firmly established equivalency of avian NCL to mammalian PFC, both functionally (Nieder, 2017) and through its macro anatomy (Güntürkün & Bugnyar, 2016), also holds for its LFP dynamics. This expands our knowledge about how higher cognition (WM) arises in birds, i.e. following the same oscillatory dynamics observed in mammals.

### Gamma modulation related to WM

Remarkably, the telencephalic LFP power dynamics in the gamma frequency range is observed across species in a similar fashion: it was elevated during stimulus encoding, contained information about stimulus location, reduced during the early delay, and ramping up towards the end of the delay (Kornblith et al., 2016; Lundqvist et al., 2016).

The observation that gamma oscillations have similar cognitive correlates in crows as in mammals, despite key anatomical differences, could point towards a key functional advantage of rhythmic population activity. This argument was previously made based on the conserved temporal properties across vastly different mammalian brain sizes (Buzsáki et al., 2013). Cortical gamma is thought to implement a winner-take-all algorithm (Fig. 4C) that simultaneously promotes selective neuronal activity without runaway excitation due to divisive normalization (Fries, 2015; Lundqvist et al., 2010). Together with analysis from single-neuron activity in crows (Hahn et al., 2021), it suggests that crow gamma could have a similar role in selection and normalization despite being implemented on a different neural substrate.

We also report that avian gamma is ‘bursty’ rather than a continuous and prolonged increase in power. The smooth elevation during stimulus encoding and the smooth increase during the end of the delay was visible only in the trial averages, at the single-trial level it was only elevated above baseline in brief bursts. Such bursts of gamma have also been observed in human and non-human primate cortex (Kucewicz et al., 2017; Lundqvist et al., 2016; Lundqvist, Herman, Warden, et al., 2018). They provide support for models in which WM information is retained by a combination of spiking and synaptic mechanisms (Fig. 4E; (Lundqvist et al., 2011; Mongillo et al., 2008; Sandberg et al., 2003)). The role of the bursts may be to facilitate reliable synaptic transmission (Lisman, 1997) and to leave a plastic synaptic mark of WM at the synapse (Miller et al., 2018). This, and other related findings, have motivated models of WM in which retention can be achieved by ‘activity silent’ mechanisms, i.e., synaptic plasticity following bursts of spiking (Lundqvist, Herman, & Miller, 2018; Miller et al., 2018; Sreenivasan & D’Esposito, 2019). However, there is an ongoing debate over these models and the more classical model of WM retention through observable sustained spiking (Constantinidis et al., 2018; Wang, 2021).

In addition to gamma oscillations, we also observed lower frequency oscillations (4-25 Hz). Similar to alpha/beta oscillations in primates, these largely showed the opposite behavior as the gamma oscillations over time (elevated when gamma was suppressed and vice versa). In cortical networks, alpha/beta oscillations are thought to play an inhibitory role and suppress gamma and the associated processing of sensory information (Händel et al., 2011; Jensen & Mazaheri, 2010; Lundqvist et al., 2016). Gamma band activity, in contrast, is associated with active encoding and decoding of WM information, e.g., when information has to enter WM, or when it is retrieved (Lundqvist et al., 2016; Roux et al., 2012; Sederberg et al., 2003). Thus, during these gamma active phases, the neuronal networks are plastic. Alpha/beta band activity is associated with retention (e.g., during the delay) that safeguards encoded information against perturbation. Our data are largely in line with these ideas, although we also observed some deviations from such mammalian data and model-predictions as outlined above.

### Deviations from mammalian models

Despite these striking similarities in the overall modulation of oscillatory activity by task epochs between birds and mammals, we also observed key deviations, in particular for load-dependent effects: despite gamma increasing during WM-encoding (load 1 vs load 0), it subsequently decreased with load. This is in stark contrast to studies from human and non-human primates in which gamma increases monotonically with load (Howard et al., 2003; Kornblith et al., 2016; Lundqvist et al., 2016; Meltzer et al., 2008; Roux et al., 2012).

From a modeling perspective, this pattern could potentially be explained by an increase of simultaneously active populations as load increases. Each population codes for distinct items. Due to the lack of columnar alignment, they could potentially cancel out each other’s contribution to the measured field when more than one is active (in contrast to the cortical alignment, Fig. 4D). However, the positive correlation between load and gamma at the end of the delay and in the choice period could speak against an anatomical explanation for this cross-species discrepancy. It should also be noted that single-neuron spiking only showed a load-dependent effect towards the end of the delay (where it increased with load, similar to mammals), suggesting there are cross-species differences in the population activity, particularly at encoding and not only in the measured LFP. This poses a challenge to existing models of working memory that tend to assume increased cognitive load is supported by increased (or at least not decreasing) population activity (Lundqvist et al., 2011). Another possible explanation could be that the birds processed the memory items differently during the sample and at the end of the delay. Because memory items were presented simultaneously, the birds might have processed them as one during the sample, but then shifted to an individual representation during the delay, like cycling through the individual colors one by one. Task-dependent changes, depending on the behavioral relevance, in the neural representations of WM items, have been reported in monkeys (Panichello & Buschman, 2021). If there’s a difference between those modes, it might explain why our observations are congruent with those of monkeys from a full sequential version of the task only at the end of the delay (Lundqvist et al., 2016).

We cannot exclude the possibility that some methodological differences (in comparison to monkeys) could have caused our observed deviations. We trained our birds to retain head fixation without restraining them which might have caused effort-related signals that attenuated some effects. Similarly, we did not explicitly control for eye movements. Importantly though, these differences were necessary to attain recordings that would allow our novel LFP analysis of purely task-related cognition. Motor-related activity in particular would have hindered such isolated analysis. Overall, the complex pattern with different load effects during encoding and choice, and non-monotonic changes from load 0 to load 3, points towards intriguing differences in the evolved implementations between mammals and birds. In addition, while gamma and alpha/beta tended to be elevated and suppressed in different parts of the trials, this relationship did not seem as strong as that in primates. For instance, the load effects for gamma and beta bursts went in the same, not opposite, directions as one would expect if they were anti-correlated.

The fact that birds have similar WM capacity, and striking similarities in the neural WM activity, makes these differences more relevant as clues towards what dynamical features are vital to support higher order cognition. Future modelling and avian neurophysiological studies hold significant promise to reveal such principles.

## Methods

Our animals, experimental setup, behavioral protocol, recording setup, and surgical procedures were previously described in (Hahn et al., 2021).

### Subjects

We worked with two hand-raised carrion crows (*Corvus corone*), held under identical housing and food protocols as described in (Hahn et al., 2021). All experimental procedures and housing conditions were carried out in accordance with the National Institutes of Health Guide for Care and Use of Laboratory Animals and were authorized by the national authority (LANUV).

### Experimental setup

Our setup consisted of an operant training chamber outfitted with a touchscreen (22’’, ELO 2200 L APR, Elo Touch Solutions Inc., CA) and an automatic feeder delivering food reward upon correct pecks on the touchscreen. We used two computer vision cameras (‘Pixy’, CMUcam5, Charmed Labs, Tx) to track the birds’ head position via a mount of two lightweight 3D-printed LEDs that was removed after each experimental session. Head-location was acquired at 50 Hz and data was smoothed by integrating over 2 frames in Matlab using custom programs on a control PC. The behavioral paradigm was executed by custom code written in Matlab using the Psychophysics (Brainard, 1997) and Biopsychology toolboxes (Rose et al., 2008). Further details about the experimental setup have been reported in (Hahn et al., 2021).

### Behavioral protocol

We trained the birds to perform a delayed non-match to sample task, previously used to test the performance under different working memory loads in primates (Buschman et al., 2011). The protocol has previously been reported by Hahn et al., (2021). Trials started with the presentation of a red dot centered on the touchscreen (for a maximum of 40 s). Centering of the head in front of the red dot for 160 ms caused the red dot to disappear and a stimulus array of two to five colored squares to appear (Fig. 1A, ‘sample’). The sample was presented for 800 ms, while the animals had to maintain head fixation and center their gaze on the screen (‘hold gaze’, no more than 2 cm horizontal or vertical displacement, and no more than 20° horizontal or vertical rotation). Failure to retain head fixation resulted in an aborted trial. The sample phase was followed by a memory delay of 1000 ms after which the stimulus array reappeared with one color exchanged. The animal indicated which of the colors had changed by pecking the respective square. Correct responses were rewarded probabilistically (BEO special pellets, in 55 % of correct trials, additional 2 s illumination of the food receptacle in 100 % of correct trials). Incorrect responses to colors that had not changed or a failure to respond within 4 s resulted in a brief screen flash and a 10 s timeout. Individual trials were separated by a 2 s inter-trial interval.

The colored squares were presented at six fixed locations on the screen (1 – 6, Fig. 1A). In each session, one pair of colors was assigned to each of the six locations. Each location had its own distinct pair. These pairs were randomly chosen from a pool of 14 colors (two color combinations were excluded since the animals did not discriminate them equally well during a pre-training). Fig. 1A gives an example. The color-change occurs for the middle right where blue (B) is presented during the sample and green (G) during the choice. In this particular session, the middle-right location could thus show either of the following colors during the sample and choice: B-G (shown in Fig. 1A); G-B; G-G; B-B; None-None. On the next session, a new random pair of colors were displayed at this location. The order of presentation of colors within a pair, the target location (where the color change occurred), and the number of stimuli in the array (two to five) were randomized and balanced across trials so that each condition had an equal likelihood to appear. The width of the colored squares was 10 degrees of visual angle (DVA) and squares were placed on the horizontal meridian of the screen and at 45.8 DVA above or below the meridian at a distance of 54 and 55.4 DVA from the center. The binocular visual field of crows is 37.6 DVA (Troscianko et al., 2012). With our arrangement on screen, combined with the head tracking, we ensured that all stimuli appeared only outside of this binocular range.

### Surgery

The surgery protocol was identical to the one reported by Hahn et al., (2021). Both animals were chronically implanted with a lightweight head-post to attach a small LED holder during the experiments. Before surgery, animals were deeply anesthetized with ketamine (50 mg/kg) and xylazine (5 mg/kg). Once deeply anesthetized, animals were placed in a stereotaxic frame. After attaching the small head-post with dental acrylic, a microdrive with a multi-channel microelectrode was stereotactically implanted at the craniotomy (Neuronexus Technologies Inc., Ann Arbor MI, DDrive). The electrode was positioned in NCL (AP 5.0, ML 13.0) of the left hemisphere (coordinates for the region based on histological studies on the localization of NCL in crows (Veit & Nieder, 2013). After the surgery, the crows received analgesics.

### Electrophysiological recordings of single-cell activity and LFP

Recordings of neuronal activity (local field potentials and single-cell spiking) were performed using chronically implanted multi-channel microelectrodes. The distance between individual recording sites (electrodes) was 50 μm. The signal was amplified, filtered, and digitized using Intan RHD2000 headstages and a USB-Interface board (Intan Technologies LLC, Los Angeles CA). The system also recorded digital event codes that were sent from the behavioral control PC using a custom IO-device (details available at www.jonasrose.net). Before each recording session, the electrodes were advanced manually using the microdrive. Recordings were started 20 minutes after the advancement, and each recording site was manually checked for neuronal signals (cellular discharges observable on an audio monitor). Signals of analysis of LFP were recorded at a sampling rate of 30 kHz and filtered with a band-pass filter at recording (1 Hz - 7.5 kHz). LFP signals were then further processed by offline down-sampling to 1 kHz. For analysis, we chose to systematically sub-sample a quarter of all electrodes used (i.e., analyzing signals from every fourth electrode, thereby achieving a reduced overlap of signal with 200 μm distance between electrodes). To verify our results we applied analysis to a second, independent subsample of the electrodes. Qualitative results from this second subsample were comparable. Data of single-cell neuronal activity for analysis of the spiking rate of the neuronal population (SFig. 5), was obtained from our previous study (Hahn et al., 2021).

### Processing of LFP results

Prior to extracting frequency power from our signals, we removed possible spike-related traces from the LFP signals using the algorithm of (Banaie Boroujeni et al., 2020). We further processed our LFP signals using the FieldTrip open-source software package for Matlab (Oostenveld et al., 2010). We extracted frequency power from the signals using Morlet-wavelet convolution with a Morlet family of 99 frequencies (2-100 Hz), with seven wavelet cycles. We screened all trials for unique trial artifacts centered around 50 Hz during processing. On rare occasions electrodes had individual trials that showed magnitudes of frequency power up to three magnitudes of power larger than the next biggest power value, we handled such artifacts by restricting data analysis to the 99^th^ percentile of power values on any electrode (i.e., excluding trials from analysis whose power values fell into the top 1 % of observed values). During manual curation of results we nonetheless observed a few electrodes with power levels exceeding their average levels at distinct time points over all frequencies (i.e. power surges not restricted to any specific frequency). Those electrodes (n = 31) were subsequently removed from data analysis altogether.

### Statistical testing of power during the trial against baseline power

We tested frequency power during the trial (in load conditions 1-3, at a 1 ms time resolution, across all individual frequencies) against baseline frequency power using a dependent samples t-statistic (i.e., testing the trial phase for a specific load against its baseline during the preceding ITI) using a permutation approach implemented in the FieldTrip toolbox. The method compares the observed t-statistic of the data (i.e., trial vs. baseline) to a null-distribution t-statistic of the permutated dataset. We used 1000 permutations, an alpha level of 5 % to determine significance, and an extreme distribution of statistical values to correct for multiple comparisons (i.e., correction was achieved by comparing observed statistical values against the most extreme (minimal and maximal) permutated values).

### Calculating gamma modulation of individual electrodes

We determined if an electrode was ‘gamma modulated’ by performing the statistical testing described above for the average power of the ‘low gamma band’ (30-59 Hz) at load 1, in 100 ms bins with 100 ms steps for the interval beginning at sample start until delay end. We classified electrodes as gamma modulated if two consecutive, non-overlapping bins had been classified as significant.

### Statistical testing of power at different loads

We tested the average change in power per added item in five frequency bands (3-7 Hz ‘theta’, 8-12 Hz ‘alpha’, 13-19 Hz ‘beta’, 30-59 Hz ‘low gamma’, and 83-98 Hz ‘high gamma’) in bins of 100 ms with a step size of 100 ms. To do so we first calculated the average power within each frequency band and bin then normalized the average power of each electrode relative to its load 1 condition (i.e., so that power at load 1 was 1 and powers at load 2 and 3 were relative to that), and finally calculated the average between the differences of load1 and load 2, and load 2 and load 3 (Eq. 1).

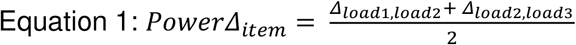

We tested if *PowerΔ*_*item*_ was significant by performing a t-test of each individual value against the null-hypothesis that it was non-different from 0, and corrected for multiple comparisons using the Bonferoni method (i.e., *α*_*crit*._ = 0.0013). We calculated the effect size of the load effect quantified by *PowerΔ*_*item*_ by performing a repeated measures ANOVA (measurements for each electrode at loads 1-3 respectively) over all electrodes and calculating the effect size (ω^2^) for all individual bins (Eq. 2).

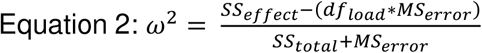

### Model comparison for location information

To investigate if LFP power contained information about the location of presented stimuli we performed a comparison of generalized linear models (GLM) applying the method of (Kornblith et al., 2016), for comparability of results. We compared a ‘full model’, containing nested load and location information, to two ‘reduced models’ where we removed location information about the ipsilateral, or contralateral locations, and replaced the respective position indicators with their sum. Each model was calculated assuming a normal distribution and its canonical ‘identity’ link function (*f*(μ) = μ). For comparison, we also assumed a gamma distribution together with its canonical link function 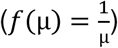. Results of both approaches were similar, but model parameters indicated that the assumption of gamma distribution did not fit all electrodes’ data, whereas the normal assumption did. We, therefore, decided to report the results of the normal models. The full model was a GLM with frequency-band power as response variable and the six possible locations as predictors. Each of the six predictors was therefore encoded as either 0 (no color at the location) or 1 (color at the location). For the reduced model we replaced three of the location indicators (either those for the contralateral locations 4-6 or those for the ipsilateral locations 1-3), by their cumulative load (i.e., 0 - 3). The reduced models thereby lacked information about the respective locations, which, if they were informative about the LFP power, would reduce the model fit (quantified by 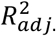). The difference between the model fits (i.e., 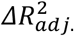) then indicates how much information was contained by the respective sides locations. We calculated this model comparison for six 400 ms bins, with a step size of 400 ms, starting 400 ms before sample onset and ending 200 ms after choice onset. We calculated if 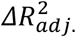 was significant in a particular bin by comparing 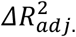 to a null-distribution 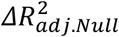 generated from the data by permutation of the data labels prior to performing the model comparison 1000 times. 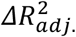 was considered significant if it was bigger than 99.17 % of permutated 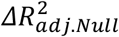 values (i.e., at an alpha level of 5 %, after Bonferoni correction for multiple comparisons).

### Calculating burst rates

Burst rates of the individual frequency bands were calculated by detecting threshold crossings of power. Frequency-band power qualifying as burst activity was defined as a power crossing a threshold of mean + 1.5*SD, for at least two consecutive cycles (periods) of the bands center frequency. For example: to classify an increase in power as a burst in the low gamma band, power had to exceed threshold levels for 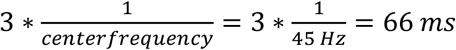. We performed this analysis with a sliding window starting at the start of the sample phase and ending with the end of the delay.

## Supplementary Material

### Supplementary section 1

**Low frequencies in Fig. 2**. Local minimum during the early sample phase occurred for a band between 8 and 20 Hz: avg. Local minimum at 8 Hz, 238 ms: 3.1780*10^6^ ± 3.4119*10^5^). The frequency band between 4 and 20 Hz had minimal power at the end of the delay (avg. local minimum within significant region at 16 Hz, 1723 ms: 2.6861×10^6 ± 3.6926*10^5).

**High frequencies in Fig. 2**. Power peaks of the higher frequencies were clustered in a band centered around 47 Hz during the middle of the sample phase (avg. local maximum of the sample phase (±standard error of the mean (SEM)) within significant region at 47 Hz, 483 ms: 1.4211*10^7^± 2.0555*10^6^) and several smaller power peaks occurred throughout the delay phase (Fig. 2).

**SFigure 1:**
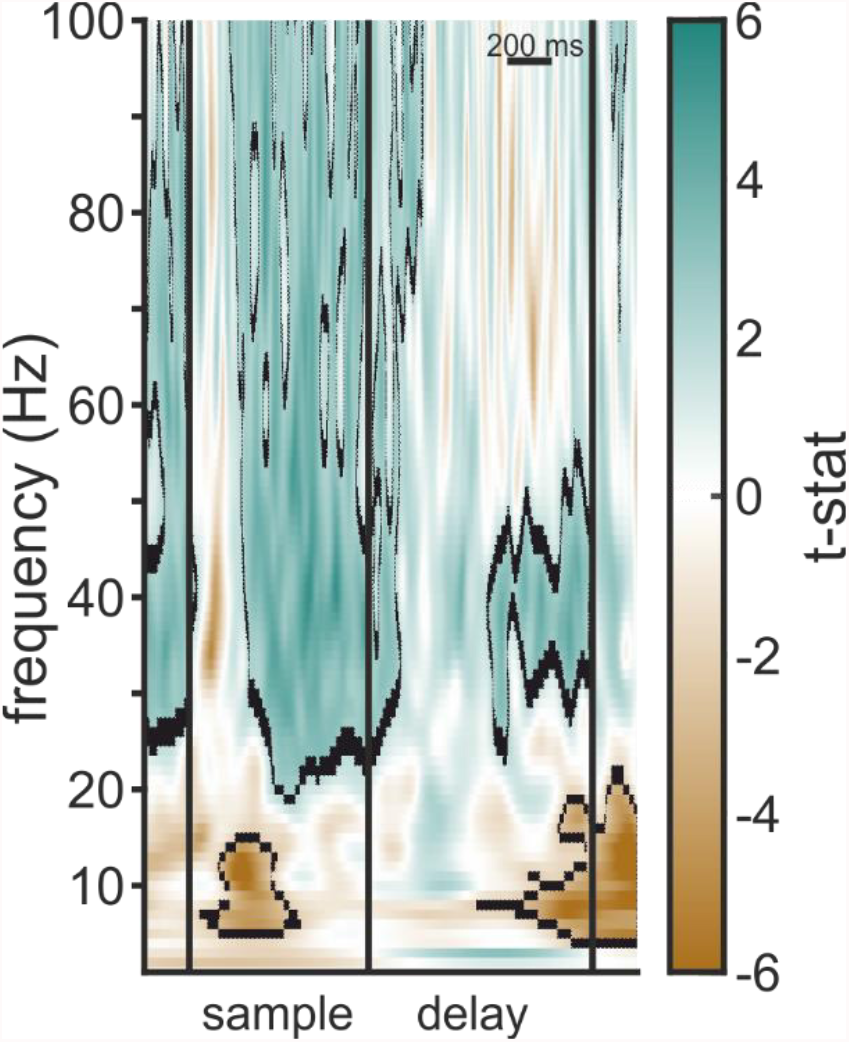
Statistical values of example electrode (Fig. 2). T-values of the significance test of load 1 vs. baseline, axes are identical to those in Fig. 2. Positive values (green) indicate that power was larger than baseline, negative values (brown) indicate that power was smaller than baseline.

### Supplementary section 2

#### Gamma power modulation throughout NCL

The peaks in power in the high gamma frequency bands were the most prominent observation at the level of individual electrodes. Was this a general effect throughout the extent of NCL or was it localized only at specific electrodes, as has been observed in monkey PFC (Lundqvist et al., 2016)? We determined if our electrodes could be classified into ‘gamma-modulated’ and ‘non-modulated’ sites, by calculating significance of a low (32-47 Hz) and high gamma (60-100 Hz) band during the sample phase for each of our electrodes (see methods for details). We found that power in the low gamma band was significantly modulated at 81.64 % of electrodes (76.72 % for high gamma). The electrodes without significant gamma modulation came from recordings obtained at locations within individual sessions (i.e., sessions in which there was no gamma modulation detected at any site), i.e., gamma modulation was either present or absent at the sites of any given recording session.

Interestingly, due to the direction of the load effect (i.e., decreasing and increasing power of the high and low-frequency bands, respectively) higher loads seemed to push power levels closer to baseline levels for both frequency bands, while an overall activation (gamma bands), or suppression (theta to beta bands) of power was present (Fig. 3A, C & E).

### Supplementary section 3

#### Power modulation based on the location of stimuli

Power of LFP contained information about the location of the colored square, mainly about the contralateral locations, but not for the ipsilateral locations. The absolute amount of information contained in LFP power in our data seemed to be slightly higher than that observed in monkeys (Kornblith et al., 2016). Position information was only present during the sample phase, not during the delay, and only for the positions contralateral to the electrode site, this indicates that gamma power plays a part in processing stimulus location unihemispherically. The optic nerve of birds is fully decussated, i.e. information observed by the right eye ends up (via the major visual pathway) exclusively in the left hemisphere (Husband & Shimizu, 2001). We designed our task to make use of this neuroanatomical isolation. Birds had to retain head fixation so that stimuli of the right side of the screen (the side contralateral to electrode implantation) were only visible to the right eye. Our results seem to reflect this manipulation indicating that interhemispheric ‘cross talk’ did not happen during WM encoding in the sample phase, suggesting independent hemispheric processing. This is in line with results from monkeys (Brincat et al., 2021; Buschman et al., 2011; Kornblith et al., 2016) and the behavioral results of this study (Balakhonov and Rose, 2017).

**SFigure 2:**
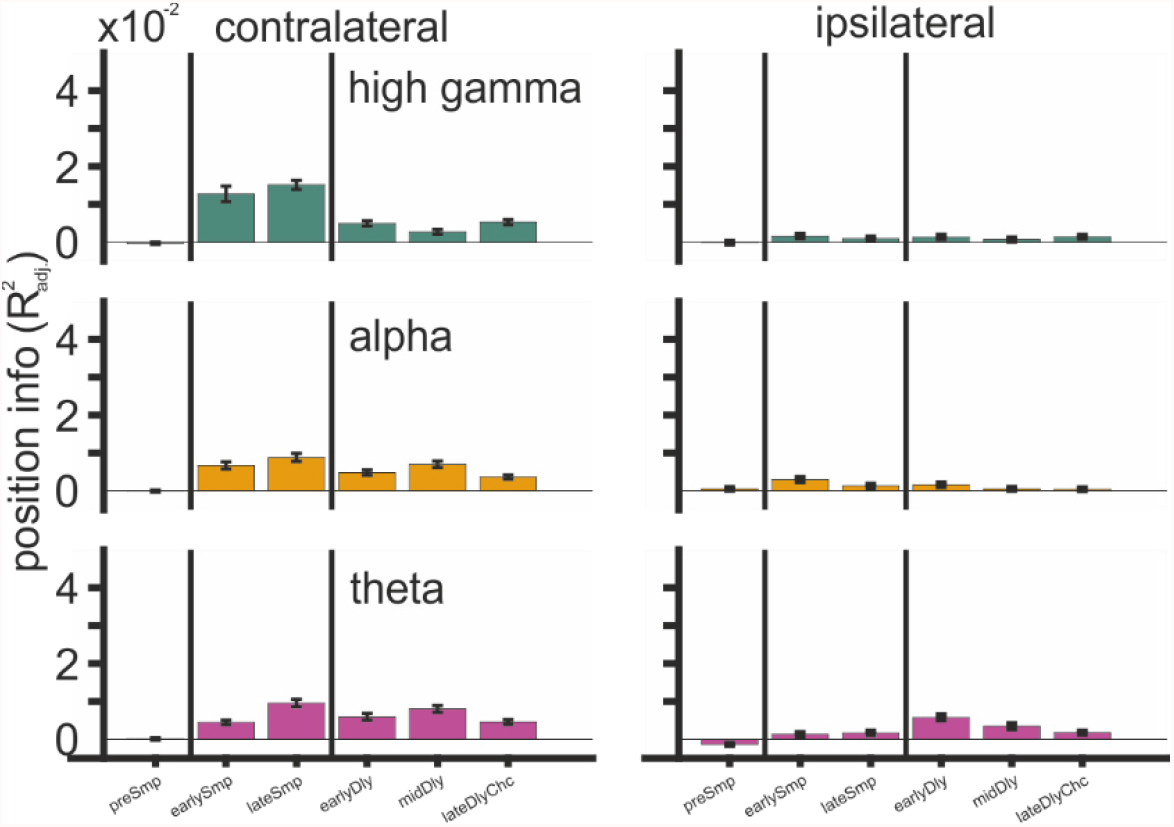
Position information 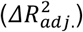 contained in average power of the theta, alpha, and high gamma bands (400 ms bins).

**SFigure 3:**
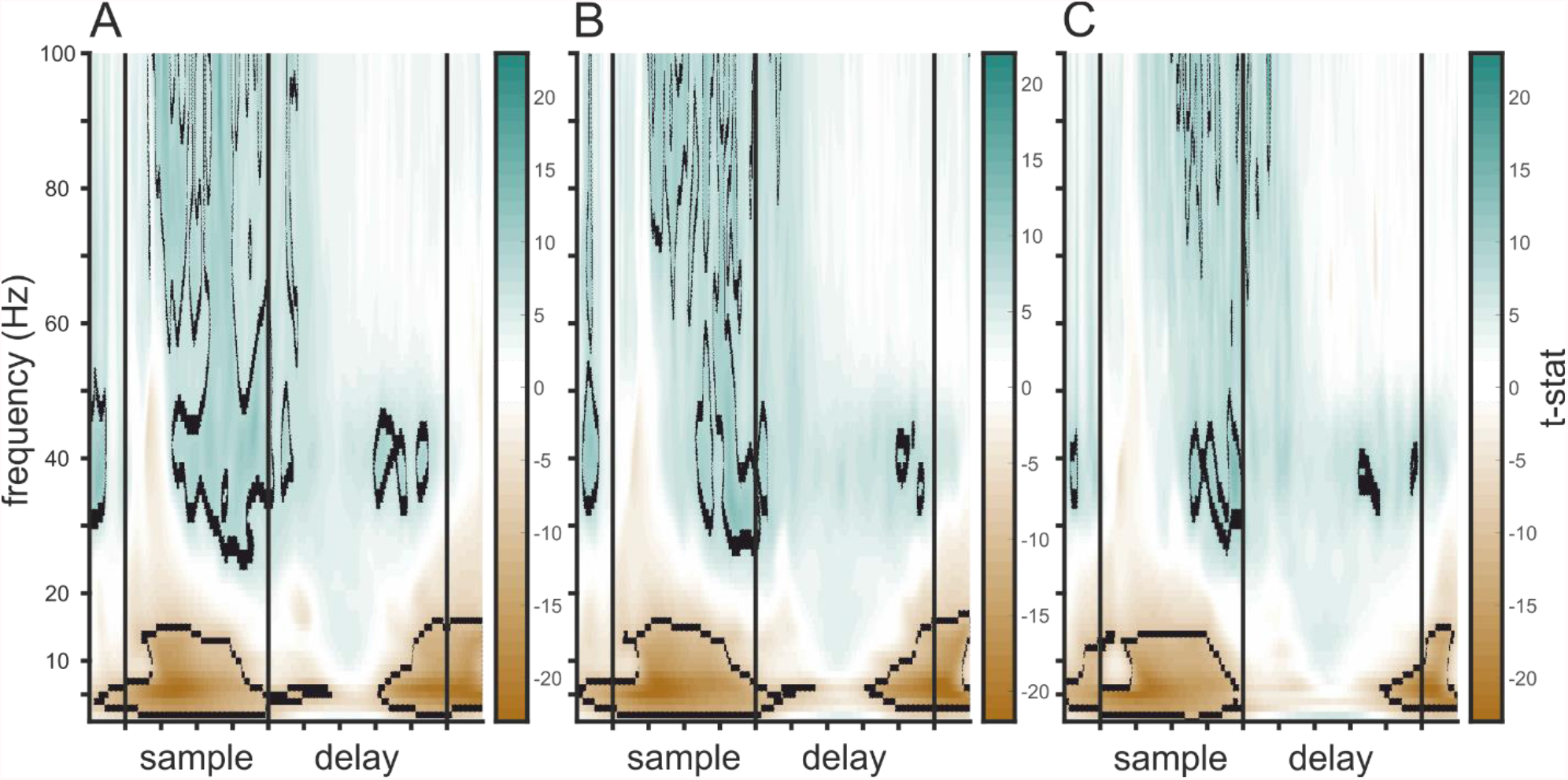
Statistical values of all significant electrodes, axes are identical to those in Fig. 3E. (A) T-values of the significance test of load 1 vs. baseline. (B) T-values of the significance test of load 2 vs. baseline. (C), T-values of the significance test of load 3 vs. baseline.

### Supplementary section 4

Bursts were also present in the alpha and high gamma frequency bands, where bursts occurred during early sample, reducing with load and remaining load-independent at a stable level during late sample, before gradually reducing throughout the delay (SFig. 4, table 4 & 5).

**SFigure 4:**
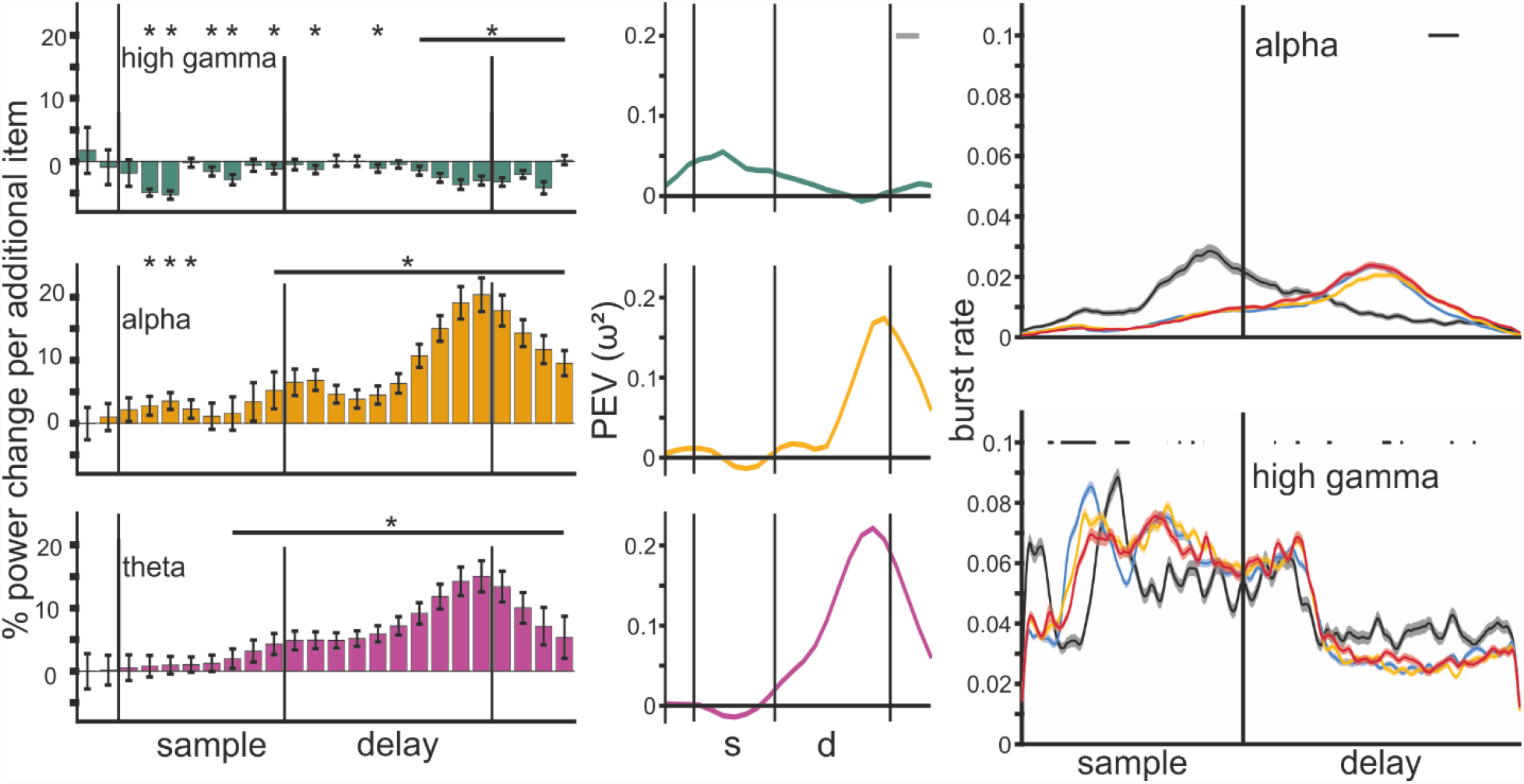
Left column: Average change in power per added item (100 ms bins) for the theta, alpha, and high gamma band. Middle: Quantification of the load effect depicted in the left column, as percent explained variance by factor power (ω^2^). Right column: Trial burst rate of alpha and high gamma frequency bands during the trial at gamma modulated sites. Black bars indicate consecutive significance between loads 1-3 (p < 0.05) over 2 cycles of the bands center frequency.

### Neuronal spiking of the population

We also compared the load effect of bursts to the spiking activity of the neurons recorded at the same time. The population of neurons increased its spiking rate during the early sample period and then gradually reduced it throughout the rest of the trial until the choice phase (SFig. 5). Load only significantly affected spiking rate towards the end of the delay, with higher loads slightly increasing spiking rate (F(1,2) = 1.3, p = 0.001, ω^2^ = 0.01, posthoc comparisons between load 1/2 and load 2/3, both p < 0.05).

**SFigure 5:**
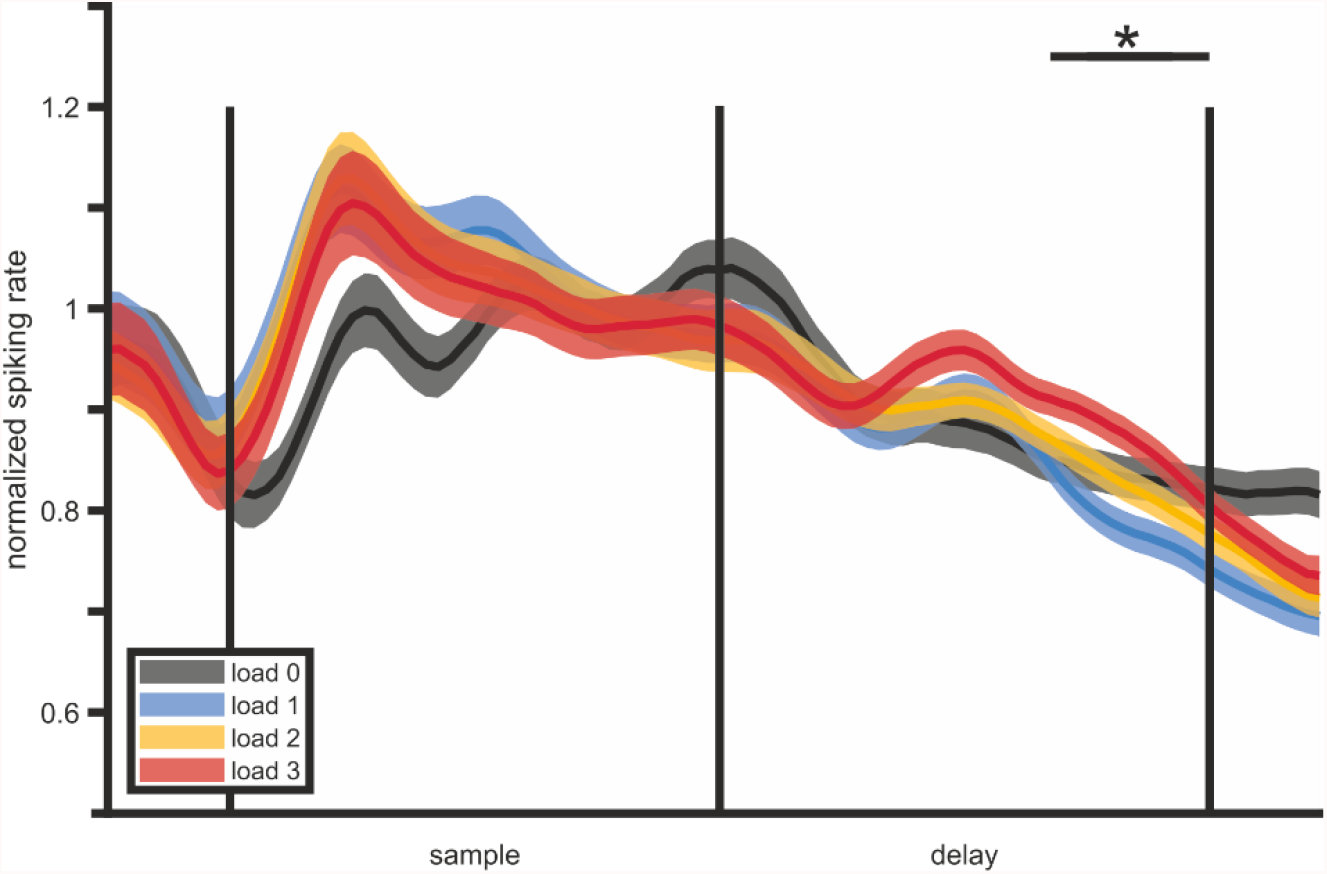
Normalized spiking rate of the neuronal population recorded during the task. The black horizontal bar indicates a significant difference between load conditions.

## Acknowledgments

We would like to thank Robert Schmidt for helpful comments on an earlier draft of the manuscript.

## Author contributions

Lukas Alexander Hahn, Data curation, Formal analysis, Investigation, Methodology, Software, Visualization, Writing - original draft, Writing - review and editing; Dmitry Balakhonov, Conceptualization, Data collection, Methodology; Mikael Lundqvist, Investigation, Visualization, Writing – review and editing; Andreas Nieder, Project administration, Resources, Writing - review and editing; Jonas Rose, Conceptualization, Funding acquisition, Methodology, Project administration, Resources, Supervision, Visualization, Writing - review and editing

